# SIK3-HDAC4-Warts Axis Functions as a Gatekeeper of Neural Stem Cell Reactivation in *Drosophila*

**DOI:** 10.64898/2026.04.11.717884

**Authors:** Yang Gao, Amanda Ng Yunn Ee, Jiaen Lin, Hongyan Wang

**Affiliations:** Neuroscience and Behavioral Disorders Programme, Duke-NUS Medical School, 8 College Road, Singapore 169857

**Author notes:** Corresponding authors: Hongyan Wang and Yang Gao, Correspondence and.

**Keywords:** Neural stem cell, reactivation, HDAC4, SIK3, Warts, SUMOylation, Hippo pathway

## Abstract

A delicate balance between the quiescent and proliferative states of neural stem cells (NSCs) is important for neurogenesis and homeostasis. Histone deacetylase 4 (HDAC4) variants are associated with neurodevelopmental disorders, however, its role in early brain development remains elusive. In this study, we demonstrate that *Drosophila* HDAC4 plays a crucial role in neural stem cells (NSCs) reactivation and brain development. Depletion of *HDAC4* results in notable defects in NSC reactivation, while its overexpression leads to premature reactivation. HDAC4 is SUMOylated at Lys902, which enhances its protein stability by preventing HDAC4 from undergoing ubiquitin-proteasome-mediated degradation. Moreover, phosphorylation of HDAC4 by salt-inducible kinase 3 (SIK3), an AMPK-related kinase, allows cytoplasmic localization of HDAC4 and enhances the association between HDAC4 and Warts, a core kinase of the Hippo pathway. This HDAC4-Wts association inhibits Warts activity, and in turn, the inactivation of the Hippo pathway, triggering NSC reactivation. Finally, genetic epistasis experiments support the SIK3-HDAC4-Warts axis during NSC reactivation. In conclusion, our findings identify HDAC4 as a molecular switch that integrates SUMOylation, ubiquitination, and the Hippo pathway to govern NSC reactivation.

## Introduction

Neural stem cells (NSCs) govern brain development, repair, nervous system regeneration and homeostasis by switching between quiescent and proliferative states. However, the majority of NSCs in the mammalian adult brain exist in the quiescent or mitotically dormant state ^1,2^. Despite this, quiescent NSCs are able to re-enter the cell cycle (reactivate) to generate neurons or glia in response to certain extrinsic signals, such as the presence of nutrition, brain injury or physical activity ^3^. This process is reversible in both aging and the absence of nutrition ^3–5^. A delicate balance between NSC quiescence and proliferation is essential for proper neurogenesis, long-term maintenance of stem cell pools, and the neurogenic capacity of the aging brain ^6–8^. Dysregulation of NSC reactivation impacts brain regeneration, repair, and overall neuroplasticity, resulting in its association with many neurodegenerative diseases, such as Alzheimer’s disease, Parkinson’s disease, Huntington’s disease, and Amyotrophic Lateral Sclerosis (ALS). As such, NSC therapy has been used as a viable treatment option in some of these neurodegenerative diseases, albeit to a limited effect ^9–16^. Variants of human orthologs of several genes that regulate NSC reactivation in *Drosophila,* such as IGF1-R, Cullin4B, and SETD8, are associated with neurodevelopmental disorders^17–25^. Therefore, understanding the molecular mechanisms underlying NSC reactivation may provide important insights into brain development and facilitate the development of therapeutics against these disorders.

*Drosophila* NSCs, also known as neuroblasts, have emerged as a powerful model system to study the mechanisms underlying NSC quiescence and reactivation *in vivo* ^7,26^. At the end of embryogenesis, *Drosophila* NSCs in the central nervous system shrink in size and enter quiescence. These NSCs will exit quiescence and reactivate in response to dietary amino acids, typically within 24 hours after larval hatching (h ALH) ^27–29^. The required nutritional signals originate from the *Drosophila* fat body which is functionally equivalent to the mammalian liver and adipose tissue ^5^. This leads to the activation of the insulin/insulin-like growth factor (IGF) signaling pathway ^30,31^ and inactivation of the Hippo signaling pathway ^32^. The blood–brain barrier (BBB) glial cells then secrete insulin-like peptides (dILPs), which diffuse directly to the underlying NSCs and reactivate them through activation of the insulin/PI3K/Akt pathway ^30,31,33^. Alternately, the inter-cellular transmembrane proteins Echinoid and Crumbs are down-regulated in the presence of nutrition, leading to the inhibition of the evolutionarily-conserved Hippo pathway that is responsible for maintaining NSC quiescence in *Drosophila* larval brains ^32,34^.

The Hippo pathway is very conserved between mammals and *Drosophila*. The core Hippo pathway consists of a kinase cascade in which Hippo (Hpo)/Mst1/2 phosphorylates Warts (Wts)/Lats1/2. This culminates at the oncogenic transcriptional coactivator Yorkie (Yki)/Yap, which is phosphorylated by upstream kinases resulting in its cytoplasmic retention ^35^. When the Hippo pathway is inactive, non-phosphorylated Yki enters the nucleus and binds to a transcription factor to enhance the expression of target genes, such as cell cycle regulators *cyclinE* (*cycE*) and the pro-proliferative and anti-apoptotic microRNA *bantam* ^35–37^. The Hippo pathway is known to maintain NSC quiescence in *Drosophila*, and this correlates with Yki localization in the cytoplasm during NSC quiescence and re-localization to the nucleus during reactivation ^32^. Furthermore, the Hippo pathway is also known to regulate NSC activity in mice. For example, Yap1 promotes adult hippocampal neural stem cell activation in mice, while its dysregulation led to the repression of hippocampal neurogenesis, aberrant cell differentiation, and partial acquisition of a glioblastoma stem cell-like signature ^38^. Wts is also a central component of the Hippo pathway, functioning as the main target for post-translational modification (PTM) during NSC reactivation. Besides its phosphorylation and activation by Hpo kinase, Wts also can be inhibited and degraded by ubiquitination and SUMOylation ^39,40^. A CRL4 E3 ubiquitin ligase composed of Cullin4, DDB1, Roc1, and a substrate receptor named Mahjong was reported to promote Wts ubiquitination and degradation, thus promoting NSC reactivation ^40^. We recently reported that the SUMO pathway is activated during NSC reactivation, promoting the SUMOylation of Wts. This modification suppresses Wts phosphorylation and kinase activity, leading to the inhibition of Hippo pathway and triggering reactivation of NSCs ^39^.

Histone deacetylases (HDACs), a family of epigenetic regulators, play a crucial role in various biological processes, including cell proliferation, differentiation, and development, by forming complexes with diverse transcription factors and co-regulators ^41^. HDACs have been shown to influence neuronal development, survival and function, and are implicated in the regulation of synaptic plasticity, as well as learning and memory ^42–44^. Additionally, HDACs affect the self-renewal and differentiation of NSCs by modulating the activity of key downstream target genes such as REST (RE1 silencing transcription factor) ^45,46^. In proliferating NSCs, transcription factors recruit HDACs to the promoters of target genes, where they repress the expression of cell cycle inhibitors like p21 and Pten, as well as neuronal-specific genes, including NeuroD, Neurogenin 1 (Ngn1) and Math1. This repression maintains NSC proliferation and self-renewal while preventing neuronal differentiation ^45,47,48^. Dysregulation of HDAC activity is associated with a wide range of brain disorders, and treatment with various HDAC inhibitors has emerged as a promising therapeutic strategy for correcting these deficiencies in neurodegenerative disease ^49^.

HDACs are classified into 4 classes based on sequence homology: class I (HDAC1, 2, 3 and 8 in vertebrates, HDAC1 and 3 in *Drosophila*), class II (IIa: HDAC4, 5, 7 and 9 in vertebrates, HDAC4 in *Drosophila*; IIb: HDAC6 and 10 in vertebrates, HDAC6 in *Drosophila*), class III (SIRT1, 2, 3, 4, 5, 6 and 7 in vertebrates, Sirt1, 2, 4, 6 and 7 in *Drosophila*), and class IV (HDAC11 in both vertebrates and Drosophila) ^50^ (Supplementary Table 1). We were drawn to the function of class IIa HDACs due to their high expression in the brain, particularly in regions of the forebrain such as the hippocampus, cortex, and amygdala ^51^. In *Drosophila*, HDAC4 is the sole class IIa Histone deacetylase, which was reported to be highly expressed in neurons and the mushroom body ^52,53^. The *Drosophila* HDAC4 (dHDAC4) protein features a long N-terminal domain and a highly conserved C-terminal catalytic domain ^54^. It shares 57% amino acid identity and 84% similarity to human HDAC4 (hHDAC4) in the deacetylase domain-containing C terminus, and 35% identity and 59% similarity across the entire protein ^52^.

HDAC4 shuttles between the nucleus and cytoplasm in both vertebrates and *Drosophila*. Alterations in the levels of nuclear and/or cytoplasmic HDAC4 controls a transcriptional program essential for synaptic plasticity and memory^41,55^, and increased abundance of nuclear HDAC4 impairs cognitive function, neuronal development and long-term memory ^52^. In vertebrates, aberrant expression or subcellular distribution of HDAC4 is linked to both neurodevelopmental and neurodegenerative disorders by affecting neuronal death and neurogenesis ^56^. These include conditions such as the Chromosome 2q37 deletion syndrome, which is characterized by developmental delay, autistic features and intellectual disability ^57–59^, as well as Alzheimer’s disease ^50,59–62^, autism ^59,63^, Huntington’s disease ^64^, stroke ^56^ and ataxia telangiectasia ^50,61,65^. In the *Drosophila* brain, dysregulation of HDAC4 results in defects in both neuronal development and the formation of long-term memories ^66^. These findings suggest that HDAC4 is neuroprotective and is a crucial positive regulator of learning and memory. Despite this, the role of HDAC4 specifically in NSCs is not clear.

In this study, we demonstrate a novel and essential role for HDAC4 in NSC reactivation and brain development. The SUMO pathway stabilizes HDAC4 protein to promote NSC reactivation. SIK3 kinase promotes HDAC4 phosphorylation and cytoplasmic accumulation to promote HDAC4 interaction with Wts kinase at its phosphorylation site and kinase domain containing C-terminal half. This interaction suppresses Wts phosphorylation and kinase activity, resulting in nuclear accumulation of Yki, therefore promoting NSC reactivation. Analysis of single-cell RNA-sequencing datasets from *Drosophila* larval brains and human fetal brains indicate that the functions of SIK3 and HDAC4 in the nervous system are evolutionarily conserved between flies and humans. Given the high conservation of the Hippo signaling pathway across species, these findings provide a potential molecular framework for understanding how dysregulation of HDAC4 contributes to neurodegenerative diseases and related neurological disorders, thereby potentially aiding the future development of therapeutic strategies targeting diseases with aberrant HDAC4 regulation.

## Results

### HDAC4 expression levels increase in NSCs during reactivation

Class IIa HDACs are reported to be highly expressed in the rat brain ^51^. In *Drosophila*, HDAC4 is the sole class IIa Histone deacetylase (Supplementary Table 1). Analysis of a published single cell RNA-sequencing dataset from late first instar larvae at 16 h ALH ^67^ revealed that HDAC4 is highly expressed in the *Drosophila* larval brain, particularly in neurons (Figure 1a and Supplementary Figure 1a, b). Most importantly, we observe an increase in *HDAC4* expression in NSCs during their reactivation (Fig 1a and Supplementary Figure 1b). Markers used for comparison include Rpl32 as a housekeeping gene, Dpn as an NSC marker known to have higher expression in active NSCs (aNSCs) than quiescent NSCs (qNSCs) ^68^; and ChAT as a neuronal marker. To detect HDAC4 in NSCs, we generated and verified an anti-HDAC4 antibody (Supplementary Figure 1c-j). A basal level expression of endogenous HDAC4 mainly localizes in cytoplasm (Supplementary Figure 1c, g). Specificity of this antibody was verified via loss of *HDAC4* in the brain or knock down of *HDAC4* in S2 cells, leading to a dramatic decrease of HDAC4 intensity in staining (Supplementary Figure 1c-d, g-h) or protein levels in Western blots (Supplementary Figure 1e-f, i-j). Consistent with the single cell RNA-sequencing data, immunostaining of the larval brain at 6 h ALH showed a 1.75-fold increase in HDAC4 protein intensity in aNSCs compared to qNSCs (Fig 1b, c). These results indicate that both mRNA and protein levels of *Drosophila* HDAC4 increase in NSCs upon reactivation. To explore the expression of Class IIa HDACs in humans, we analyzed a published single cell RNA-sequencing dataset obtained from human fetal brain during peak neurogenesis and early gliogenesis ^69^. The results revealed that all Class IIa HDACs are expressed across various cell types in the human fetal brain. HDAC4 is notably enriched in apical radial glia cells (aRG, human neural stem cells), HDAC5 in intermediate progenitor cells (IPCs), HDAC7 in vascular cells, and HDAC9 in Cajal-Retzius cells (CRs) which are specialized neurons essential for cortical development (Supplementary Figure 1k). Our analyses suggest that both *Drosophila* and human HDAC4 are expressed in neural stem cells.

**Figure 1.**
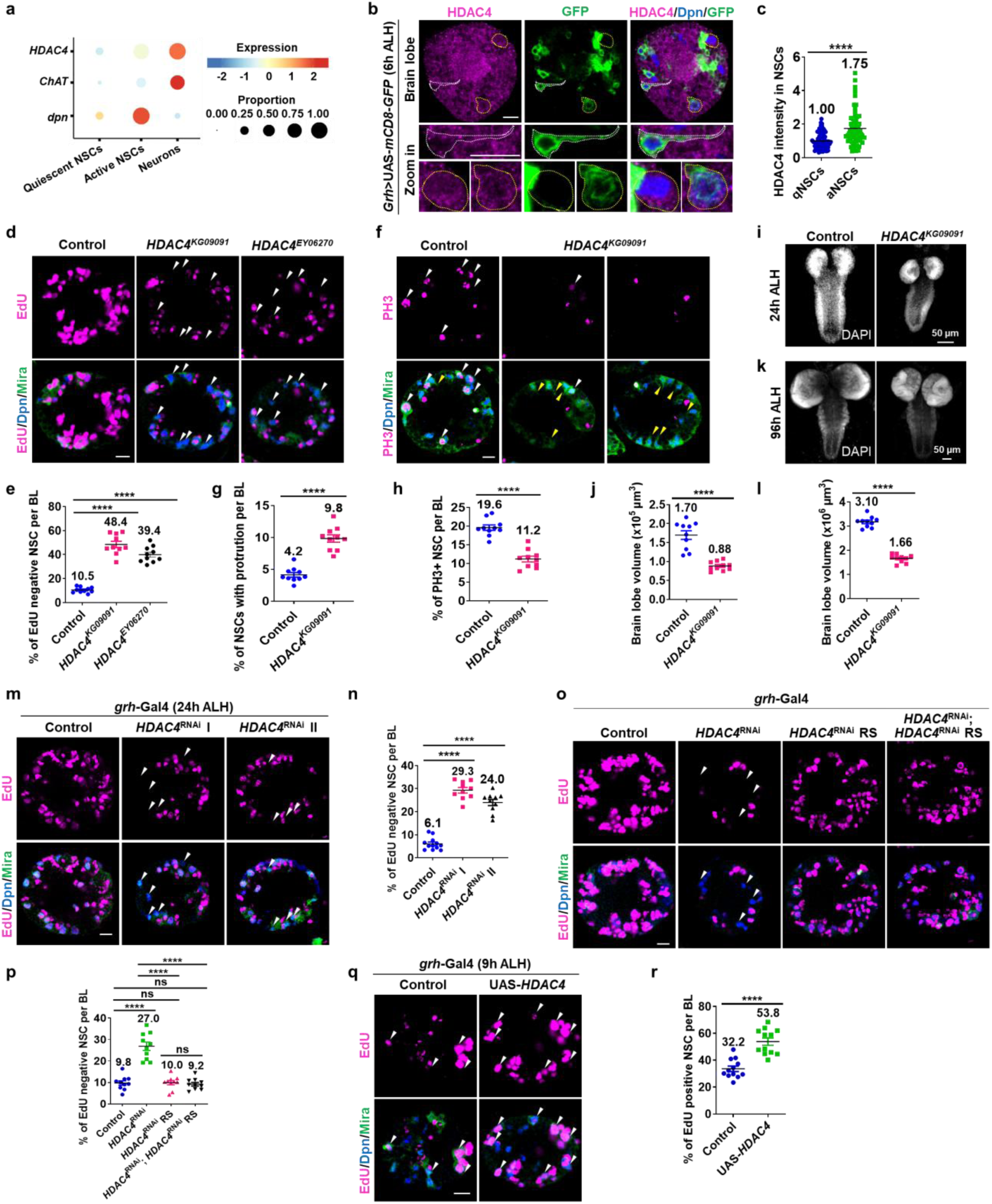
HDAC4 is necessary for NSC reactivation and brain development. **a** Analysis of a published dataset of single cell RNA-sequencing obtained from late first instar larvae at 16 h ALH ^67^. *dpn* is an NSC marker as a positive control; ChAT is a neuron marker. **b** At 6 h ALH, larval brains express mCD8-GFP driven by *grh*-GAL4. White dotted lines mark the quiescent NSCs (qNSCs). Yellow dotted lines mark the active NSCs (aNSCs). **c** Quantification of HDAC4 intensity (normalized to GFP) at 6 h ALH for (**b**) in aNSCs (big Dpn+ cells) (1.75 ± 1.03, P = 1.27E-07, n = 10) and qNSCs (Dpn+ cells with protrusion) (1.00 ± 0.47, n = 10). **d**, **m, o** At 24 h ALH, larval brain lobes from various genotypes were analyzed for EdU incorporation. NSCs were marked by Dpn and Mira. White arrows point to EdU negative qNSCs. **e** Quantification of EdU- NSCs per brain lobe (BL) for (**d**). Control (yw), 10.5 ± 2.3, n = 10; *HDAC4^KG09091^*, 48.4 ± 8.4, P = 5.1-11, n = 10; *HDAC4^EY06270^*, 39.4 ± 7.2, P = 4.3E-10, n = 10. **f** Larval brain lobes from control (yw) and *HDAC4^KG09091^* were labeled with Dpn, PH3 and Mira. White arrows point to PH3+ NSCs, yellow arrows point to qNSCs with protrusion. **g** Quantification of NSCs retaining cellular protrusion per BL for (**f**). Control, 4.2 ± 1.1, n = 10; *HDAC4^KG09091^*, 9.8 ± 1.8, P = 1.4E-07, n = 10. **h** Quantification of PH3+ NSCs per BL for (**f**). Control, 19.6 ± 2.3, n = 10; *HDAC4^KG09091^*, 11.2 ± 2.5, P = 2.8E-07, n = 10. **i**, **k** Maximum intensity z-projection of larval brains from control (yw) and *HDAC4^KG09091^* were stained with DAPI at 24 and 96 h ALH, respectively. **j** Quantification of brain volume in (**i**). Control, 1.70 ± 0.36, n = 10; *HDAC4^KG09091^*, 0.88 ± 0.12, P = 2.65E-06, n = 10. **l** Quantification of brain volume in (**k**). Control, 3.1 ± 0.31, n = 10; *HDAC4^KG09091^*, 1.66 ± 0.16, P = 1.70E-10, n = 10. **n** Quantification of EdU- NSCs per BL for (**m**). Control (β-gal), 6.1 ± 2.7, n = 11; *HDAC4*^RNAi^-1, 29.3 ± 3.9, P = 3.4E-13, n = 10; *HDAC4*^RNAi^-2, 24 ± 4.3, P = 1.3E-10, n = 10. **p** Quantification of EdU- NSCs per BL for (**o**). Control (β-gal), 9.8 ± 3.3, n = 10; *HDAC4*^RNAi^-II (VDRC 20522), 27 ± 6.2, P = 3.8E-07, n = 10; *HDAC4*^RNAi^ resistant (RS), 10 ± 3.1, P = 0.89, n = 10; *HDAC4*^RNAi^; *HDAC4*^RNAi^ RS, 9.2 ± 2.6, P = 1.3E-07, n = 10. **q** At 9 h ALH, larval brain lobes in control (β-galRNAi) and UAS-*HDAC4* lines driven by *grh-*Gal4 were analyzed for EdU incorporation. White arrows point to EdU+ NSCs. **r** Quantification of EdU+ NSCs per BL in (**q**). Control, 32.2 ± 7.1, n = 12; UAS-*HDAC4*, 53.8 ± 9.5, P = 5.8E-06, n = 12. Data are presented as mean ± SD. ns for P > 0.05, **** for P ≤ 0.0001. Scale bars: 10 μm.

### HDAC4 is required for NSC reactivation and brain development

To investigate the role of HDAC4 in NSC reactivation, we examined two *HDAC4* loss-of-function alleles, *HDAC4^KG09091^*and *HDAC4^EY06270^*. The *HDAC4^KG09091^* allele is a classic hypomorphic mutant with a SUPor-P transposon insertion within the HDAC4 locus, leading to reduced expression and function ^70^. Similarly, the *HDAC4^EY06270^* allele is a P-element insertion hypomorph that presumably disrupts endogenous HDAC4 expression and function. At 24 h ALH, the majority of NSCs in control larval brain were reactivated and incorporated with 5-ethynyl-2’-deoxyuridine (EdU), with only 10.5% of NSCs remaining quiescent and negative for EdU (Figure 1d, e). In contrast, the percentage of quiescent NSCs that were EdU-negative was significantly increased at 48.4% and 39.4% in *HDAC4^KG09091^* and *HDAC4^EY06270^* mutants respectively (Figure 1d, e), indicating a defect in NSC reactivation. We further assessed NSC quiescence by looking for the retention of primary cellular protrusions which is a known hallmark of quiescent NSCs ^33,71^, marked by cortical/cytoplasmic Miranda (Mira). Consistently, we found that there was a significant higher percentage of Mira-positive NSCs that still extended their cellular protrusions in *HDAC4* mutant brains (9.8%) as compared to 4.2% in control larval brains (Figure 1f, g). Furthermore, the number of mitotic NSCs indicated by positive phospho-Histone H3 (PH3) signal was significantly reduced from 19.6% in control larval brains to 11.2% in *HDAC4^KG09091^* mutant brains (Figure 1f, h). Moreover, *HDAC4^KG09091^*continued to exhibit NSC reactivation defects at 48h, 72h and 96h ALH (Supplementary Figure 1l, m), suggesting that the loss of *HDAC4* delays NSC reactivation even past the later stages of neurodevelopment. The delayed NSC reactivation corroborates with the significantly reduced brain lobe volume of *HDAC4^KG09091^* mutants at 0.88 X 10^5^ µm^3^, compared to control larval brain volume of 1.7 X 10^5^ µm^3^ (Figure 1i, j). This microcephaly-like phenotype also manifested at 96 h ALH, where *HDAC4^KG09091^* mutant brain lobes show a similar reduction in volume at 1.66 X 10^6^ µm^3^ compared to control at 3.1 X10^6^ µm^3^ (Figure 1k, l). To further validate the importance of HDAC4 in NSC reactivation, we knocked down *HDAC4* in NSCs (driven by NSC-specific driver *grainy head*-Gal4 (*grh*-Gal4)) using two independent RNAi lines at 24 h ALH. Consistent with the mutants, the percentage of EdU-negative quiescent NSCs significantly increased from 6.1% in control larval brains to 29.3% and 24% in the *HDAC4*^RNAi^ lines (Figure 1m, n), indicating a significant defect in NSC reactivation. Moreover, the NSC reactivation defects upon HDAC4 RNAi were rescued by expression of an RNAi-resistant HDAC4 transgene generated by codon modification (Figure 1o, p), confirming the specificity of the HDAC4 RNAi effect. These data indicate that HDAC4 is required for NSC reactivation and proper brain development.

### Overexpression of *HDAC4* leads to premature NSC reactivation

To further explore how HDAC4 regulates NSC reactivation, we obtained and verified the Flag-tagged *HDAC4* transgenic flies and overexpressed *HDAC4* in NSCs (Supplementary Figure 1n-p). Overexpression of *HDAC4* resulted in a dramatic increase in HDAC4 intensity, localizing in both the cytoplasm and nucleus, appearing as punctate aggregates in the latter compartment (Supplementary Figure 1n, p). At 9 h ALH, majority of the wild-type NSCs remained in a quiescent state, indicated by only 32.2 % of NSCs incorporated EdU (Figure 1q, r). Remarkably, overexpression of *HDAC4* by g*rh*-Gal4 significantly increased the percentage of proliferative NSCs marked by EdU incorporation to 53.8% (Figure 1q, r), suggesting that *HDAC4* overexpression triggers premature NSC reactivation. However, overexpression of *HDAC4* failed to promote NSC reactivation in larvae on a sucrose-only diet even at 24 h ALH (Supplementary Figure 1q, r), suggesting that HDAC4-mediated NSC cell cycle reentry is nutrient-dependent. Taken together, our results indicate that HDAC4 is both necessary and sufficient for promoting NSC reactivation under nutrient-rich conditions, and indispensable for brain development.

### SUMO pathway promotes HDAC4 SUMOylation at Lys902

We recently reported that the SUMO pathway promotes NSC reactivation ^39^, and human HDAC4 was reported to be SUMOylated at Lys-559 ^72^. Utilizing the Joined Advanced SUMOylation Site and Sim Analyser (JASSA) ^73^, a different lysine in *Drosophila* HDAC4 was predicted to be SUMOylated at Lys902 (HDAC4-K902) (Supplementary Table 2), which corresponds to the conserved human HDAC4-K737 residue (Supplementary Table 3). To confirm the validity of this prediction, we conducted SUMOylation assays in S2 cells. Cells were transiently transfected with Flag-HDAC4 (∼150 kD), either alone or in combination with Myc-tagged wild-type (WT) Smt3 or Smt3^G87A, G88A^ (denoted as Smt3^AA^, the conjugation deficient mutant of SUMO ^74,75^). Following immunoprecipitation (IP) with a Flag antibody, the resulting protein complexes were detected at the anticipated >150 kD bands corresponding to SUMOylated Flag-HDAC4 (Figure 2a). Notably, wild-type *smt3* overexpression resulted in a 1.29-fold increase in SUMOylated Flag-HDAC4 (Figure 2a, b), while overexpression of Smt3^AA^ significantly reduced HDAC4 SUMOylation by 0.65-fold (Figure 2a, b), suggesting that HDAC4 is indeed SUMOylated.

**Figure 2.**
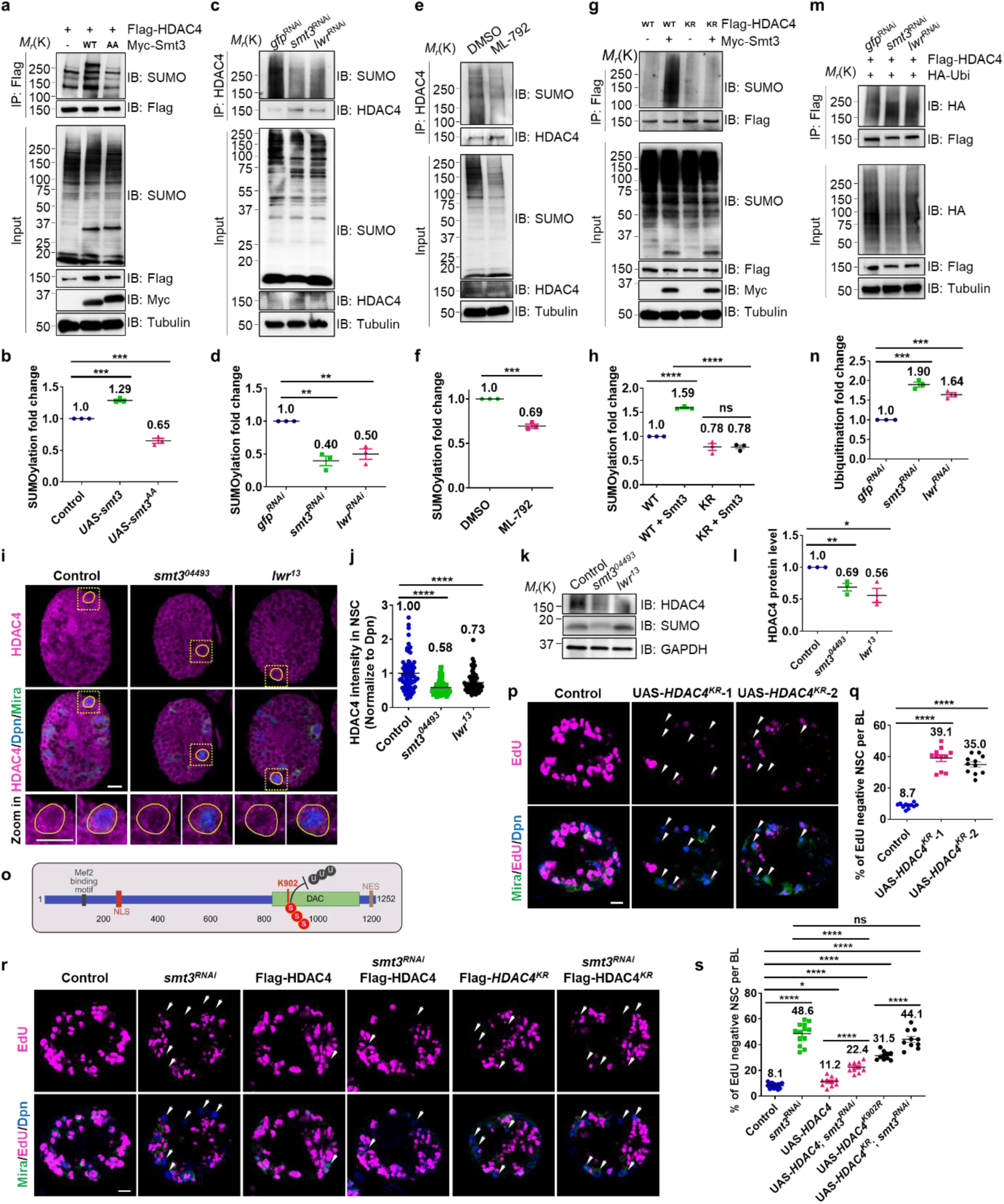
SUMO modification of HDAC4 promotes its protein stability and NSC reactivation. **a**, **c, e, g** SUMOylation assays in S2 cells. IP was conducted with anti-Flag antibody (**a, g**) or anti-HDAC4 antibody (**c, e**), and western blots were performed by using anti-SUMO and anti-Flag/HDAC4 to detect SUMOylated HDAC4 and overall levels of Flag-HDAC4/endogenous HDAC4, respectively. Tubulin serves as a loading control. **a** Flag-HDAC4 and Myc-Smt3WT or Myc-Smt3AA were co-transfected into S2 cells. **c** S2 cells were treated with double stranded RNA against gfp, smt3 or lwr for 72 h to induce gene knockdown. **e** 6 h before harvested, S2 cells were treated with DMSO or ML-792. **g** Flag-HDAC4WT or Flag-WtsKR was co-transfected with Myc-Smt3 into S2 cells. **b, d, f, h** Quantification of HDAC4 SUMOylation for (**a**, **c, e, g**) respectively, n = 3. **b** Control (Flag-Wts), 1; Flag-HDAC4 + Myc-Smt3WT, 1.29 ± 0.04, P = 0.0002; Flag-Wts + Myc-Smt3AA, 0.65 ± 0.07, P = 0.0008. **d** Control (gfpRNAi), 1; smt3RNAi, 0.4 ± 0.13, P = 0.001; lwrRNAi, 0.5 ± 0.14, P = 0.003. **f** DMSO, 1; ML-792, 0.69 ± 0.04, P = 0.002. **h** Control (Flag-HDAC4WT), 1; Flag- HDAC4WT + Myc-Smt3, 1.59 ± 0.03, P = 5.2E-06; Flag-HDAC4KR, 0.78 ± 0.12; Flag- HDAC4KR + Myc-Smt3, 0.78 ± 0.07, P = 0.98, P = 4.6E-05. **I** Larval brain lobes from control (*yw*), *smt3^04493^* and *lwr^13^* at 24 h ALH. **j** Quantification graph of HDAC4 intensity (normalized to Dpn) in NSCs in (**i**). Control, 1+0.5, n = 11; *smt3^04493^*, 0.58 ± 0.21, P = 4.1E-10, n = 10; *lwr^13^*, 0.73 ± 0.29, P = 9.2E-05, n = 10. **k** Protein extracted from larval brains of control (*yw*), *smt3^04493^*and *lwr^13^* at 24 h ALH was used for Western blot. **l** Quantification graph of HDAC4 protein levels (normalized to GAPDH) in (**k**), n=3. Control, 1; *smt3^04493^*, 0.69 ± 0.1, P = 0.007; *lwr^13^*, 0.56 ± 0.19, P = 0.02. **m** Ubiquitination assay in S2 cells, cells were treated with double stranded RNA against gfp, smt3 or lwr for 72 h to induce gene knockdown. Flag-HDAC4 and HA-Ubi were co-transfected into S2 cells. S2 cells were treated with 10 µM MG132 for 4 h before harvest. IP was conducted with anti-Flag antibody. Western blots were performed by using anti-HA and anti-Flag to detect ubiquitinated HDAC4 and overall levels of Flag-HDAC4, respectively. **n** Quantification of HDAC4 ubiquitination for (**m**). Control (gfpRNAi), 1; smt3RNAi, 1.9 ± 0.11, P = 0.0001; lwrRNAi, 1.64 ± 0.09, P = 0.0003. **o** Illustration of HDAC4 protein and HDAC4 SUMOylation site. MEF2: Mef2 binding domain; NLS: nuclear localization signal; DAC: deacetylase domain; K902: Lysine 902 residue; S: SUMO; U: ubiquitin; NES: nuclear export signal. **p** At 24 h ALH, larval brain lobes from control (β-galRNAi) and UAS-*HDAC4^KR^* lines driven by *grh*-Gal4 were analyzed for EdU incorporation. NSCs were marked by Dpn and Mira. White arrows point to EdU- qNSCs. **q** Quantification of EdU- NSCs per BL for (**p**). Control, 8.7 ± 1.6, n = 11; UAS-*HDAC4^KR^*-1, 39.1 ± 7.0, P = 2.2E-12, n = 11; UAS-*HDAC4^KR^*-2, 35 ± 5.9, P = 7.4E-11, n = 11. **r** At 24 h ALH, larval brain lobes in indicated genotypes were analyzed for EdU incorporation. NSCs were marked by Dpn and Mira. White arrows point to EdU negative quiescent NSCs. **s** Quantification graph of EdU- NSCs per BL for genotypes in (**r**). Control (β-galRNAi; β-galRNAi), 8.1 ± 2.1, n = 14; smt3RNAi; β-galRNAi, 48.6 ± 8.74 P = 1.4E-15, n = 13; β-galRNAi; UAS-HDAC4, 11.2± 0.01 P = 1.4E-11, P = 0.01, n = 10; smt3RNAi; UAS-HDAC4, 22.4 ± 4.0, P = 10E-09, P = 4.5E-06, n = 10; β-galRNAi; UAS-HDAC4KR, 31.5 ± 3.3, P = 3.9E-16, n = 10; smt3RNAi; HDAC4KR, 44.1 ± 7.3, P =1.62E-14, P = 9.1E-05, P=0.2, n = 10. Data are presented as mean ± SD. ns for P > 0.05, * for 0.05≤ P ≤ 0.01, ** for P ≤ 0.01, *** for P ≤ 0.001, **** for P ≤ 0.0001. Scale bars: 10 μm.

Subsequently, we examined the SUMOylation of endogenous HDAC4 in S2 cells following the depletion of *SUMO*/*smt3* or SUMO E2 *lwr* with dsRNA treatment. We confirmed a reduction in overall SUMOylation in *smt3*- or *lwr*- knockdown groups in the input fraction using the anti-SUMO antibody. Using the HDAC4 antibody, we observed a significant decrease in the SUMOylation of HDAC4 in the IP fraction by 0.4-fold and 0.5-fold in the *smt3*- and *lwr*-knockdown groups respectively (Figure 2c, d), indicating that both SUMO and SUMO E2 are required for HDAC4 SUMOylation. We also utilized ML-792, which is an effective inhibitor of SUMOylation acting at the SUMO activating enzyme (SAE, SUMO E1) ^76^. Upon IP of HDAC4, we observed a significant, 0.69-fold reduction of HDAC4 SUMOylation when treated with the ML-792 inhibitor compared to the DMSO control group (Figure 2e, f). Moreover, ML-792 inhibitor treatment also suppressed the overall SUMOylation in the input fraction (Figure 2e). Therefore, SUMO E1 enzyme is also indispensable for facilitating HDAC4 SUMOylation.

To determine whether HDAC4-K902 serves as the primary site for SUMOylation on HDAC4, the K902 residue in HDAC4 was replaced with an arginine (R) residue for a subsequent SUMOylation assay. In S2 cells, Myc-Smt3 overexpression led to a 1.59-fold increase in the SUMOylation of Flag-HDAC4, while the SUMOylation of Flag-HDAC4^K902R^ (HDAC4^KR^) remained unchanged (Figure 2g, h). This observation strongly suggests that K902 represents a major site of SUMOylation on HDAC4.

### SUMOylation stabilizes HDAC4 by preventing its ubiquitin-proteasome-mediated degradation

SUMOylation is a type of post-translational modification which may affect various aspects of the substrate protein, including its activity, stability, or subcellular localization ^77^. To investigate the effect of SUMO modification on HDAC4, we tested the expression of HDAC4 in the context of SUMO or SUMO E2 depletion in larva brains at 24 h ALH. In loss of function alleles of both *smt3* and *lwr*, HDAC4 fluorescence intensity in brain lobes showed an obvious decrease, with the intensity in NSCs reduced by 0.58-fold in *smt3* mutant and 0.73-fold in *lwr* mutant, respectively (Figure 2i, j). Consistently, western blotting using larval brains at 24 h ALH showed a significant reduction in HDAC4 protein levels in both *smt3* and *lwr* mutants by 0.69-fold and 0.56-fold respectively (Figure 2k, l). Knockdown of *smt3* and *lwr* via RNAi in S2 cells also showed a significant decrease of HDAC4 protein levels (Supplementary Figure 2a, b). Quantitative real-time PCR revealed no significant change in *HDAC4* mRNA levels in *smt3^04493^* mutant larval brains compared to control at 24 h ALH (Supplementary Figure 2c, d), consistent with the notion that the SUMO pathway regulates HDAC4 at the post-translational level. We further conducted cycloheximide (CHX) chase assays in S2 cells to block protein biosynthesis and detect HDAC4^WT^ or HDAC4^KR^ protein levels without or with Smt3 co-overexpression in a time-course experiment. Notably, Smt3 overexpression slowed down the degradation of HDAC4^WT^ but not HDAC4^KR^ (Supplementary Figure 2e, f), suggesting that the SUMO modification stabilizes HDAC4. Interestingly, treatment of S2 cells with the proteasome inhibitor MG132 was found to inhibit HDAC4 protein degradation caused by *smt3* or *lwr* knockdown (Supplementary Figure 2g, h), suggesting that HDAC4 protein degradation caused by SUMO or SUMO E2 depletion relies on the ubiquitin-proteasome-mediated degradation pathway.

In order to test if HDAC4 SUMOylation alters its ubiquitination levels, we performed ubiquitination assay in S2 cells. Cells were transiently co-transfected with Flag-HDAC4 and HA-Ubiquitin, along with the depletion of *smt3* or *lwr* via dsRNA treatment, while dsRNA-*gfp* was used as control. Following IP with a Flag antibody, the resulting immune complexes were detected at the anticipated >150 kD bands corresponding to ubiquitinated Flag-HDAC4 (Figure 2m), where knockdown of *smt3* and *lwr* resulted in a 1.9-fold and 1.64-fold increase in ubiquitinated Flag-HDAC4 respectively (Figure 2m, n). In contrast, overexpression of wild-type *smt3* suppresses HDAC4 ubiquitination (Supplementary Figure 2i, j; compare lane 2 to lane 1), while overexpression of the conjugation deficient *smt3* (*smt3^AA^*) significantly promotes HDAC4 ubiquitination (compare lane 3 to lane 1). These observations suggest that SUMOylation stabilizes HDAC4 by preventing its ubiquitination. Intriguingly, the ubiquitination level of HDAC4^K902R^ is much lower than wild-type HDAC4 (Supplementary Figure 2i, j; compare lane 4 to lane 1), suggesting that HDAC4-K902 is also a main target site for ubiquitination, in addition to SUMOylation. Consistent with this notion, the ubiquitination level of HDAC4^K902R^ was insensitive to co-overexpression of *smt3* or *smt3^AA^* (Supplementary Figure 2i, j; compare lanes 5-6 to lane 3), suggesting that HDAC4^K902R^ is deficient in both SUMOylation and ubiquitination. Our results are consistent with the previous reports that SUMO and ubiquitin can competitively bind to the same lysine of the substrate ^78,79^(Figure 2o). Given that the SUMO protein levels and *SUMO* mRNA levels dramatically increased with NSC reactivation ^39^, it is conceivable that the SUMO modification overrides ubiquitination to stabilize HDAC4 protein during NSC reactivation.

To further test this hypothesis, we generated two independent transgenic fly lines expressing HDAC4 SUMOylation/ubiquitination-deficient HDAC4^K902R^ (Venus-tagged) and validated their expression by immunostaining (Supplementary Figure 2k). Overexpression of either *HDAC4^K902R^* line using *grh*-Gal4 at 24 h ALH resulted in pronounced NSC reactivation defects, where EdU incorporation assays revealed that 39.1% and 35% of *HDAC4^K902R^*-overexpressing NSCs failed to incorporate EdU, compared with 8.7% in control brains (Figure 2p, q). These results suggest that HDAC4^K902R^ might act as a dominant-negative form. Moreover, overexpression of wild-type HDAC4 significantly suppressed the delayed reactivation phenotype caused by *smt3* depletion (Figure 2r, s; *smt3*^RNAi^ = 48.6%. UAS-*HDAC4*; *smt3*^RNAi^ = 22.4%), despite wild-type HDAC4 overexpression alone inducing only mild NSC reactivation defects (Figure 2r, s; UAS-*HDAC4* = 11.2%, compared to control =8.1%). In contrast, overexpression of the HDAC4^K902R^ failed to rescue the NSC reactivation defects induced by *smt3* depletion (Figure 2r, s; *smt3*^RNAi^ = 48.6% vs UAS-*HDAC4^K902R^*; *smt3*^RNAi^ = 44.1%). These findings suggest that SUMOylation and ubiquitination of HDAC4 modulate its function during NSC reactivation.

### HDAC4 interacts with C-terminal Wts containing phosphorylation sites and kinase domain

Several publications in the past few years reported potential correlation between HDACs and the Hippo pathway in both mammals and *Drosophila* ^80–84^. For example, HDAC inhibitors are known to suppress the primary target of the Hippo pathway YAP ^83,84^. Mammalian *HDAC3* depletion is also known to lead to transcriptional changes enriched in Hippo pathway-related genes ^81,82^. In *Drosophila*, *HDAC1* depletion reduces Yki signaling ^80^, indicating direct involvement of HDAC1 in the Hippo pathway. Since the Hippo pathway is known to be one of the most important regulators of NSC reactivation, exploring the interaction between HDAC4 and the Hippo pathway would likely reveal key and meaningful insights into the mechanism behind NSC reactivation.

Mouse HDAC3 regulates iron homeostasis in the liver through the Hippo pathway, while *Drosophila* HDAC1 inhibits apoptosis via inhibiting the Hippo pathway ^80,81,82^. Given that Hippo pathway inhibits NSC reactivation/proliferation in *Drosophila* ^32,39^ and in mammals ^38^, we firstly explored the potential interaction between *Drosophila* HDAC4 and the Hippo pathway. Wts is a key kinase in the Hippo pathway that integrates upstream kinases signals and transmits them to the downstream primary effector Yki. Interestingly, co-Immunoprecipitation (co-IP) assays conducted in S2 cells between HDAC4 and Wts in either direction of IP showed a physical association (Figure 3a, b). To determine the specific region of Wts responsible for its interaction with HDAC4, we performed co-IP assays between HDAC4 and truncated forms of Wts: N-terminal half containing the PPxY motifs (Wts-NT) and the C-terminal half containing the kinase domain and phosphorylation sites (Wts-CT) (Figure 3c). The co-IP results demonstrated that HDAC4 interacts with Wts-CT, but not with Wts-NT (Figure 3d, e). These findings suggest that the interaction between Wts and HDAC4 occurs specifically within the C-terminal region of Wts that encompasses its phosphorylation sites and kinase domain. Given that the C-terminal region of Wts is critical for its phosphorylation by Hippo kinase and for its kinase activity toward the downstream effector Yki, the interaction between HDAC4 and Wts may have important regulatory implications for Hippo pathway activity.

**Figure 3.**
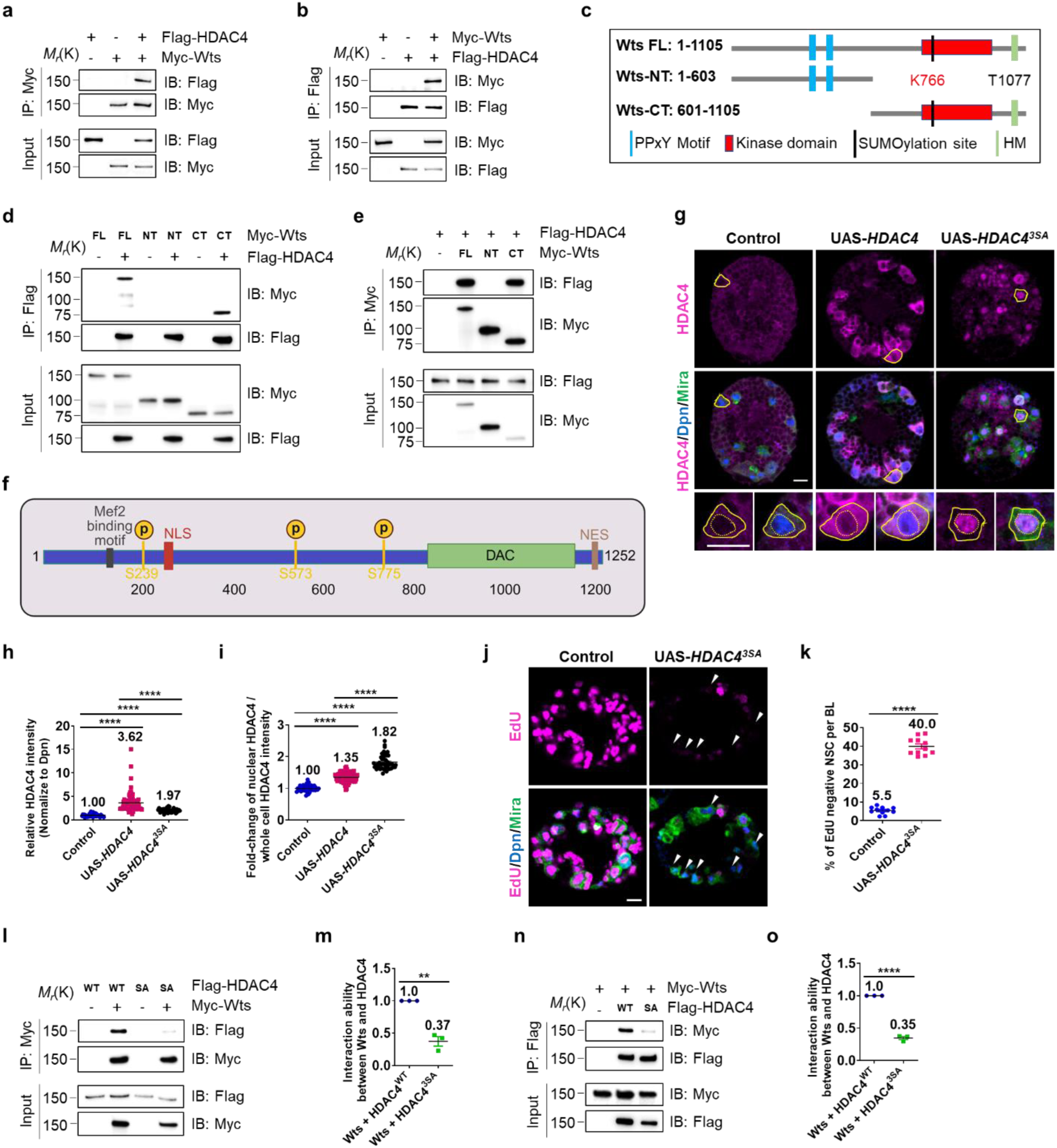
HDAC4 interacts with C-terminal Wts containing phosphorylation sites and kinase domain. **a**, **b** In S2 cells, Co-IP of exogenously expressed Myc-Wts with Flag-HDAC4 and vice versa. **c** The illustration of Wts protein. PPxY motif is required for Yki interaction. The SUMOylation site K766 is within the kinase domain of Wts protein. HM is hydrophobic motif, T1077 is the Thr phosphorylated by Hpo, and this site is within the hydrophobic motif of Wts protein. Wts-NT: 1-603 aa, Wts-CT: 601-1105 aa. **d, e** In S2 cells, Co-IP of exogenously expressed Myc-Wts FL, NT and CT with Flag-HDAC4 and vice versa. **f** Illustration of HDAC4 protein and HDAC4 phosphorylation sites. P: phosphorylation. **g** Larval brain lobes in control (β-galRNAi), UAS-*HDAC4* and UAS-*HDAC4^3SA^* driven by *grh-*Gal4 at 24 h ALH. **h** Quantification graph of HDAC4 intensity (normalized to Dpn) in NSCs in (**g**). Control, 1, n = 11; UAS-*HDAC4*, 3.62 ± 1.4, P = 8.5E-14, n = 11; UAS-*HDAC4^3SA^*, 1.97 ± 1.4, P = 8.5E-14, n = 11. **i** Quantification graph of fold-change of nuclear HDAC4/whole cell HDAC4 intensity in NSCs in (**g**). Control, 1, n = 11; UAS-*HDAC4*, 1.35 ± 1.4, P = 8.5E-14, n = 11; UAS-*HDAC4^3SA^*, 1.82 ± 1.4, P = 8.5E-14, n = 11. **J** At 24 h ALH, larval brain lobes in control (β-galRNAi) and UAS-*HDAC43SA* lines driven by *grh-*Gal4 were analyzed for EdU incorporation. White arrows point to EdU- NSCs. **k** Quantification of EdU- NSCs per BL in (**j**). Control, 5.46 ± 4.6, n = 10; UAS-*HDAC43SA*, 40 ± 11.3, P = 2.3E- 07, n = 10. **l, n** In S2 cells, Co-IP of exogenously expressed Myc-Wts with Flag-HDAC4 WT/3SA and vice versa. **m, o** Quantification graph of fold-change of interaction ability between Wts and HDAC4 WT/3SA in (**l, n**), respectively. Data are presented as mean ± SD. ** for P ≤ 0.01, **** for P ≤ 0.0001. Scale bars: 10 μm.

### Phosphorylated HDAC4 localizes to the cytoplasm and associates with Wts to promote NSC reactivation

HDAC4 is known to shuttle between the nucleus and cytoplasm in both vertebrates and *Drosophila*. Dysregulation of nuclear-cytoplasmic shuttling of HDAC4 is associated with several neurodevelopmental and neurodegenerative disorders ^56^. As in hHDAC4, dHDAC4 also contains a nuclear localization signal (NLS) ^85^, a nuclear export sequence (NES) ^59^ and a transcription factor MEF2 binding region, which is indispensable for HDAC4 nuclear localization ^53,85,86^ (Figure 3e). In addition, the cytoplasmic localization of HDAC4 has been reported to be regulated via both salt-inducible kinase 3 (SIK3) and calcium/calmodulin-dependent protein kinase (CaMK) mediated phosphorylation of conserved serine residues (S246, S467 and S623 in hHDAC4, S239, S573 and S775 in dHDAC4), which allows binding of the cytoplasmic chaperone 14-3-3ζ ^85,87–92^(Figure 3f). Conversely, dephosphorylation of HDAC4 by the serine/threonine phosphatase PP2A allows its re-entry into nucleus ^93^.

A previous study with *Drosophila* HDAC4^3SA^ shows an upregulation of nuclear HDAC4 in the mushroom body, impaired neuronal development, long-term memory and eye development ^52,53^. We thus wondered if expressing HDAC4^3SA^ in NSCs would also result in a similar nuclear accumulation of HDAC4 in NSCs that subsequently impairs NSC reactivation. At 24 h ALH, while overexpression of either wild-type HDAC4 or HDAC4^3SA^ in NSCs showed an expected increase in HDAC4 intensity (Figure 3g, h), wild-type HDAC4 mainly localized in the cytoplasm of NSCs, while HDAC4^3SA^ accumulated in the nuclei of NSCs (Figure 3g, i). Furthermore, EdU incorporation assays showed severe NSC reactivation defects upon overexpression of HDAC4^3SA^ (percentage of EdU negative NSCs; control = 5.5%, UAS-*HDAC4^3SA^* = 40%, Figure 3j, k), suggesting elevated nuclear HDAC4 impairs NSC reactivation.

Interestingly, co-IP assays between Wts and HDAC4 or HDAC4^3SA^ revealed that HDAC4^3SA^ markedly disrupted the interaction with Wts. Quantitative analysis showed that the interaction between Wts and HDAC4^3SA^ is reduced to less than 40% of that observed between Wts and wild-type HDAC4 (Figure 3l-o). To further validate the association between HDAC4 mutations and Wts, we employed proximity ligation assay (PLA), a technique that enables the detection of protein–protein interactions with high specificity and sensitivity ^94^. In S2 cells, co-overexpression of Wts with wild-type HDAC4 or HDAC4^K902R^ showed strong PLA signal, but co-overexpression of HDAC4^3SA^ and Wts showed dramatically reduced PLA signal (Supplementary Figure 3a-c). Indeed, HDAC4^3SA^ showed nuclear accumulation and strong aggregation in S2 cells (Supplementary Figure 3a). These results suggested that HDAC4 phosphorylation results in its cytoplasmic localization, allowing it to associate with Wts during NSC reactivation.

### *Sik3* is highly expressed in brains of both *Drosophila* and humans

SIK3, a well-established kinase, is known to phosphorylate and localize HDAC4 to the cytoplasm in *Drosophila* and mammals ^87,95,96^. We thus wondered if this mechanism functions similarly in the NSCs of the larval brain. We first examined the expression pattern of *Sik3* in the fly larval brain by analyzing a published single cell RNA-sequencing dataset obtained from late first instar larvae at 16 h ALH ^67^. Similar to *HDAC4*, *Sik3* is highly expressed in fly larval brain, with particularly strong expression in neurons (Figure 4a and Supplementary Figure 4a). Notably, *Sik3* expression in NSCs is also elevated during NSC reactivation, displaying a trend comparable to that observed for *HDAC4* (Figure 4a). We also explored the expression of human *SIK3* by analyzing a published single-cell RNA-sequencing dataset obtained from human fetal brain during peak neurogenesis and early gliogenesis ^69^. *SIK3* exhibited an expression pattern similar to that of *HDAC4*, with both genes showing high expression in apical radial glia (aRG), the NSCs in humans (Supplementary Figure 4b). These findings suggest that *Sik3* is expressed in NSCs in both *Drosophila* and human brains.

**Figure 4.**
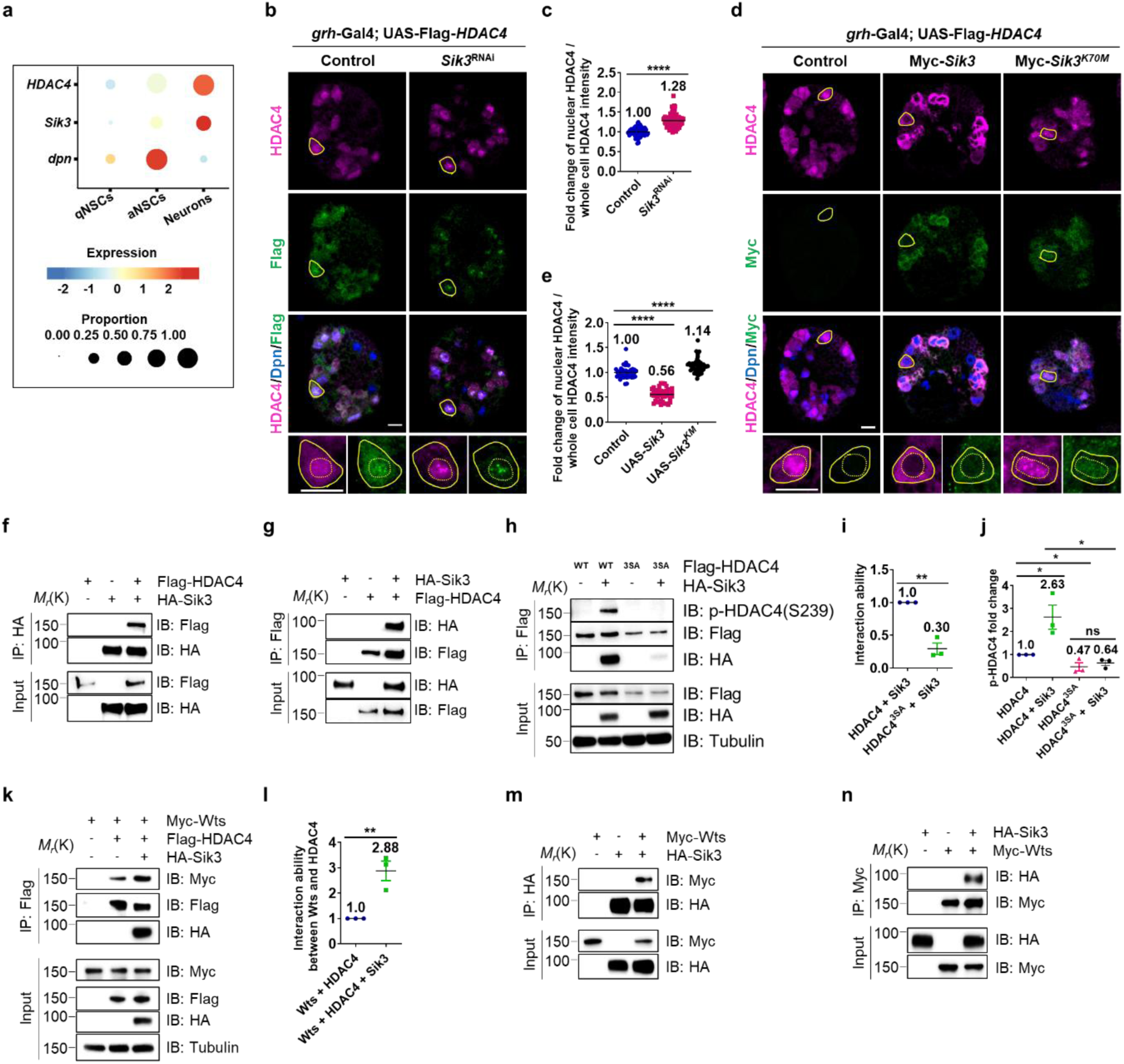
SIK3 promotes HDAC4 cytoplasmic localization in a kinase activity–dependent manner. **a** Re-analysis of a published dataset of single cell RNA-sequencing obtained from late first instar larvae at 16 h ALH^67^ for the expression of *Sik3* and *HDAC4* in quiescent NSCs, active NSCs and neurons. **b** Larval brain lobes in control (β-galRNAi; UAS-Flag-HDAC4) and *Sik3*^RNAi^; UAS-Flag-HDAC4 driven by *grh-*Gal4 at 24 h ALH. **c** Quantification graph of fold-change of nuclear HDAC4/whole cell HDAC4 intensity in NSCs in (**b**). Control, 1±0.54, n = 10; *Sik3*^RNAi^; UAS-Flag-HDAC4, 1.28 ± 0.17, P = 2.2E-29, n = 12. **d** Larval brain lobes in control (β-galRNAi; UAS-Flag-HDAC4), UAS-Myc-*Sik3*; UAS-Flag-HDAC4 and UAS-Myc-*Sik3^K70M^*; UAS-Flag-HDAC4 driven by *grh-*Gal4 at 24 h ALH. **e** Quantification graph of fold-change of nuclear HDAC4/whole cell HDAC4 intensity in NSCs in (**b**). Control, 1±0.12, n = 10; UAS-Myc-*Sik3*; UAS-Flag-HDAC4, 0.56 ± 0.12, P = 2.8E-29, n = 10. UAS-Myc-*Sik3^K70M^*; UAS-Flag-HDAC4, 1.14 ± 0.14, P = 3.5E-06, n = 10. **f**, **g** In S2 cells, Co-IP of exogenously expressed HA-Sik3 with Flag-HDAC4 and vice versa. **h** In S2 cells, Co-IP of exogenously expressed HA-Sik3 with Flag-HDAC4 WT/3SA. The phosphorylated HDAC4 at Ser239 was detected in IP group. **i** Quantification graph of fold-change of interaction ability between SIK3 and HDAC4 WT/3SA in (**h**), n=3. HDAC4 + Sik3, 1; HDAC4^3SA^ + Sik3, 0.3 ± 1.4, P = 0.001. **j** Quantification graph of p-HDAC4-S239 fold-change in (**h**), n=3. HDAC4, 1; HDAC4 + Sik3, 2.63 ± 0.91, P = 0.04; HDAC4^3SA^, 0.47 ± 0.31, P = 0.04; HDAC4^3SA^ + Sik3, 0.64 ± 0.2, P = 0.04. P = 0.02, P = 0.48. **k** In S2 cells, Co-IP of exogenously expressed Myc-Wts with Flag-HDAC4 without or with HA-Sik3 co-expressing. **l** Quantification graph of fold-change of interaction ability between Wts and HDAC4 in (**k**), n=3. Wts + HDAC4, 1; Wts + HDAC4 + Sik3, 2.88 ± 0.67, P = 0.008. **m**, **n** In S2 cells, Co-IP of exogenously expressed HA-Sik3 with Myc-Wts and vice versa. Data are presented as mean ± SD. * for 0.05≤ P ≤ 0.01, ** for P ≤ 0.01, **** for P ≤ 0.0001. Scale bars: 10 μm.

### SIK3 regulates cytoplasmic localization of HDAC4 in a kinase activity–dependent manner

Next, we investigated whether SIK3 regulates the subcellular localization of HDAC4 in NSCs. At 24 h ALH, Flag-tagged HDAC4, detected by both anti-HDAC4 and anti-Flag antibodies, was distributed relatively evenly between the cytoplasm and nucleus in control brain lobes. In contrast, knockdown of *Sik3* resulted in pronounced nuclear accumulation of HDAC4, with a notable increase in HDAC4 aggregation within the nucleus, accompanied by a dramatic reduction of cytoplasmic HDAC4 (Figure 4b, c). Conversely, overexpression of wild-type Myc-tagged *Sik3* in NSCs promoted the translocation of HDAC4 from the nucleus to the cytoplasm (Figure 4d, e). However, overexpression of a kinase-dead *Sik3* (*Sik3^K70M^*) in NSCs resulted in nuclear accumulation and aggregation of HDAC4 (Figure 4d, e). Wild-type SIK3 co-localized with HDAC4 in the cytoplasm, whereas SIK3^K70M^ co-translocated with HDAC4 and co-localized within the nucleus (Figure 4d, e).

Consistent with the *in vivo* results, knockdown or overexpression of *Sik3* in S2 cells produced similar effects. In S2 cells, depletion of *Sik3* using two independent dsRNA led to significant nuclear accumulation of HDAC4 compared with the *gfp*-RNAi control (Supplementary Figure 4c-e). Conversely, overexpression of HA-tagged wild-type *Sik3* promoted the translocation of HDAC4 from the nucleus to the cytoplasm (Supplementary Figure 4f, g). However, overexpression of the kinase-dead mutant *Sik3^K70M^* induced nuclear accumulation and aggregation of HDAC4 (Supplementary Figure 4f, g), resembling the effects observed upon expression of HDAC4^3SA^ in NSCs and S2 cells (Figure 3f and Supplementary Figure 3a). Collectively, these results indicate that Sik3 is required for the cytoplasmic localization of HDAC4 in a kinase activity–dependent manner.

We next investigated whether SIK3 promotes HDAC4 cytoplasmic localization through direct interaction. In S2 cells, co-IP assays using either HDAC4 or SIK3 as bait revealed robust interaction between the two proteins (Figure 4f, g). Notably, the interaction between SIK3 and the phosphorylation-deficient mutant HDAC4^3SA^ was markedly reduced at 30% of the interaction observed with wild-type HDAC4 (Figure 4h, i). Moreover, co-expression of *Sik3* markedly enhanced the phosphorylation of wild-type HDAC4 by 2.63-fold, whereas it had no detectable effect on the phosphorylation of HDAC4^3SA^ (Figure 4h, j). These results demonstrate that SIK3 interacts with and phosphorylates HDAC4 in a manner that depends on the three serine residues.

### SIK3 interacts with Wts and promotes the interaction between Wts and HDAC4

Because SIK3 phosphorylates HDAC4 at S246, S467 and S623, HDAC4^3SA^ is considered a phosphorylation–defective mutant of HDAC4 ^92^. Given that HDAC4^3SA^ interacts poorly with Wts, we next wondered if SIK3 modulates the HDAC4–Wts interaction. Co-IP assays revealed that co-expression of *Sik3* significantly enhanced the interaction between HDAC4 and Wts by 2.88-fold (Figure 4k, l), suggesting that SIK3-mediated phosphorylation of HDAC4 enhances the interaction between HDAC4 and Wts. Since *Drosophila* SIK3 was reported as a negative regulator of the Hippo signalling pathway in fly wings ^97^, we proceeded to further explore if SIK3 interacts with Wts. Interestingly, in S2 cells, co-IP assays between SIK3 and Wts revealed a clear interaction between the two proteins (Figure 4m, n). Taken together, these findings indicate that SIK3 interacts with both Wts and HDAC4, and this interaction enhances the association between Wts and HDAC4.

### *Sik3* loss-of-function results in microcephaly and defects in NSC reactivation

We next explored the effect of SIK3 on NSC reactivation. We examined two potential *Sik3* loss-of-function alleles, *Sik3^f03280^* and *Sik3^EY06260^*. The *Sik3^f03280^* is a PiggyBac transposon insertion allele within the *Sik3* genomic locus, and *Sik3^EY06260^* allele is a P-element insertion mutant where the EPgy2 transposon has inserted into the *Sik3* locus ^92^. At 24 h ALH, the percentage of EdU-negative quiescent NSCs significantly increased from 10.5% in control to 30.4% and 44.9% in *Sik3^f03280^* and *Sik3^EY06260^* mutants respectively (Figure 5a, b). Brain lobe volumes of *Sik3^EY06260^* mutants at 24 h ALH was significantly reduced to 0.79 X 10^5^ µm^3^, compared with 1.47 X 10^5^ µm^3^ in control larval brains (Figure 5c, d). This microcephaly-like phenotype persisted at 96 h ALH, with *Sik3^EY06260^* mutant brain lobes exhibiting a similar reduced volume of 1.97 X 10^6^ µm^3^, compared with 2.73 X10^6^ µm^3^ in control (Figure 5e, f). These results indicate that SIK3 plays an important role in NSC reactivation and brain development. Furthermore, knockdown of *Sik3* in NSCs using three independent RNAi lines resulted in significant NSC reactivation defects (Figure 5g, h).

**Figure 5.**
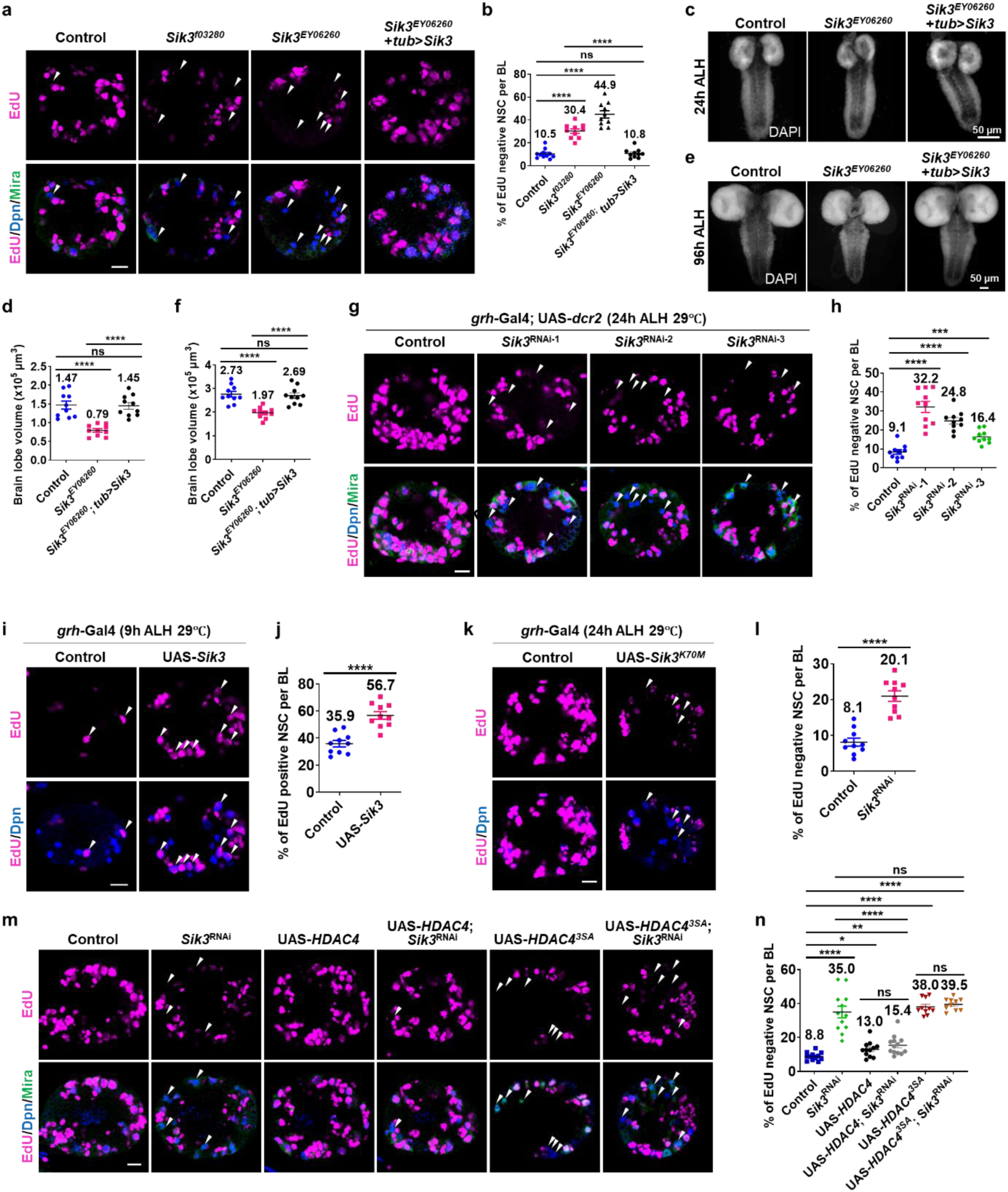
SIK3 is indispensable for NSC reactivation and brain development. **a**, **g, k, m** At 24 h ALH, larval brain lobes from various genotypes were analyzed for EdU incorporation. NSCs were marked by Dpn and Mira. White arrows point to EdU negative qNSCs. **b** Quantification of EdU- NSCs per brain lobe (BL) for (**a**). Control (yw), 10.5 ± 4.1, n = 11; *Sik3^f03280^*, 30.4 ± 6.3, P = 5.4E-08, n = 10; *Sik3^EY06260^*, 44.9 ± 10.2, P = 2.9E-09, n = 10. *Sik3^EY06260^*; *tub*>*Sik3* 10.8 ± 4.0, P = 0.8, P = 1.1E-08, n = 10. **c**, **e** Maximum intensity z-projection of larval brains from control (yw) and *Sik3^EY06260^* were stained with DAPI at 24 and 96 h ALH, respectively. **d** Quantification of brain volume in (**c**). Control, 1.47 ± 0.34, n = 10; *Sik3^EY06260^*, 0.79 ± 0.15, P = 1.8E-05, n = 10; *Sik3^EY06260^*; *tub*>*Sik3* 1.45 ± 0.28, P =0.89, P = 3.5E-6, n = 10. **f** Quantification of brain volume in (**e**). Control, 2.73 ± 0.4, n = 10; *Sik3^EY06260^*, 1.97 ± 0.23, P = 6.6E-05, n = 10; *Sik3^EY06260^*; *tub*>*Sik3* 2.69 ± 0.37, P = 0.86, P = 5.6E-05, n = 10. **h** Quantification of EdU- NSCs per brain lobe (BL) for (**g**). Control (β-galRNAi), 8.4 ± 3.8, n = 10; *Sik3*^RNAi^-1, 32.2 ± 9.3, P = 6.3E-07, n = 10; *Sik3*^RNAi^-2, 24.8 ± 5.3, P = 2.6E-07, n = 10. *Sik3*^RNAi^-3, 16.4 ± 3.4, P = 0.0001, n = 10. **i** At 9 h ALH, larval brain lobes in control (β-galRNAi) and UAS-*Sik3* lines driven by *grh-*Gal4 were analyzed for EdU incorporation. White arrows point to EdU+ NSCs. **j** Quantification of EdU+ NSCs per BL in (**i**). Control, 35.9 ± 7.5, n = 10; UAS-*Sik3*, 56.7 ± 8.9, P = 2.2E-05, n = 10. **l** Quantification of EdU- NSCs per BL in (**k**). Control (β-galRNAi), 8.1 ± 3.5, n = 10; UAS-*Sik3^K70M^*, 21.0± 4.6, P =1.5E-06, n = 10. **n** Quantification of EdU- NSCs per BL in (**m**). Control (β-galRNAi; β-galRNAi), 8.8 ± 2.4, n = 12; β-galRNAi; *Sik3*RNAi, 35 ± 11.8, P = 1.5E-07, n = 12; UAS-HDAC4; β-galRNAi, 13 ± 4.5, P = 0.01, n = 12; UAS-HDAC4; *Sik3*RNAi, 15.4 ± 5.6, P = 0.001, P = 1.8E-05, P = 0.2, n = 13; UAS-HDAC43SA; β-galRNAi, 38± 4.7, P = 3.5E-14, n = 10; UAS-HDAC43SA; *Sik3*RNAi, 39.5 ± 3.5, P = 2.5E-16, P = 0.26, P = 0.44, n = 10. Data are presented as mean ± SD. ns for P > 0.05, * for 0.05≤ P ≤ 0.01, **** for P ≤ 0.0001. Scale bars: 10 μm.

Conversely, overexpression of Myc-tagged *Sik3* in NSCs at 9 h ALH promoted premature NSC reactivation, with the percentage of EdU positive activated NSCs increasing significantly from 35.9% in control brains to 56.7% in *Sik3*-overexpression brains (Figure 5i, j). However, overexpression of the kinase-dead mutant *Sik3^K70M^* in NSCs at 24 h ALH led to pronounced NSC reactivation defects, with the percentage of EdU negative NSCs increasing significantly from 8.1% in control brains to 20.1% in *Sik3^K70M^* overexpression brains (Figure 5k, l), indicating that SIK3 promotes NSC reactivation in a kinase activity–dependent manner. Furthermore, overexpression of *Sik3* driven by *tubulin*-Gal4 (*tub*-Gal4) fully rescued NSC reactivation defects (Figure 5a, b) and microcephaly-like phenotypes (Figure 5c-f) in *Sik3^EY06260^* mutants, further supporting the role of SIK3 in brain development and NSC reactivation.

We next investigated if *HDAC4* overexpression could rescue the reactivation defects caused by *Sik3* depletion. Overexpression of wild-type *HDAC4* in NSCs significantly suppressed the NSC reactivation defects induced by *Sik3* knockdown, as evidenced by a reduction in the percentage of EdU-negative NSCs from 35% in *Sik3*-RNAi brains to 15.4% in UAS-*HDAC4*; *Sik3*-RNAi brains (Figure 5m, n). In contrast, overexpression of the phosphorylation-deficient mutant *HDAC4^3SA^* failed to rescue this phenotype (*Sik3*-RNAi: 35%; UAS-*HDAC4^3SA^*; *Sik3*-RNAi: 39.5%. Figure 5m, n). Collectively, these results demonstrate that Sik3 is required for HDAC4 phosphorylation and cytoplasmic localization, thereby promoting NSC reactivation and proper brain development.

### SIK3-HDAC4 axis is required for Yki nuclear localization in NSCs

Upon inactivation of the Hippo pathway, non-phosphorylated transcriptional co-activator Yki enters the nucleus and facilitates NSC reactivation. Conversely, activation of the Hippo pathway leads to Yki phosphorylation and its cytoplasmic retention, thereby maintaining NSC quiescence ^32,34^. Since both SIK3 and HDAC4 interact with Wts, we tested whether they affect the subcellular localization of the Yki. At 24 h ALH, Yki predominantly localized within the nucleus of NSCs in control larval brains. Interestingly, the percentage of nuclear Yki (nYki) dramatically reduced to 65.7% in *Sik3* mutant NSCs and 43.3% in *HDAC4* mutant NSCs as compared to around 88% in control (Figure 6a-b, d-e). Furthermore, the intensity of nYki in NSCs significantly decreased to 0.81-fold in *Sik3* mutant NSCs and 0.85-fold in *HDAC4* mutant NSCs compared to control (Figure 6a, c and d, f). These observations suggest that the Hippo pathway is active in *Sik3* or *HDAC4* depletion. Moreover, overexpression of the SIK3 phosphorylation defective *HDAC4^3SA^* in NSCs at 18 h ALH significantly reduced the proportion of NSCs with nuclear Yki, from 89.2% in control to 69.9% in *HDAC4^3SA^* -expressing brains (Figure 6g, h). This reduction was accompanied by a 0.65-fold decrease in nYki intensity compared with control (Figure 6g, i). Together, these data indicate that SIK3 and cytoplasmic HDAC4 are required for efficient nuclear localization of Yki in NSCs.

**Figure 6.**
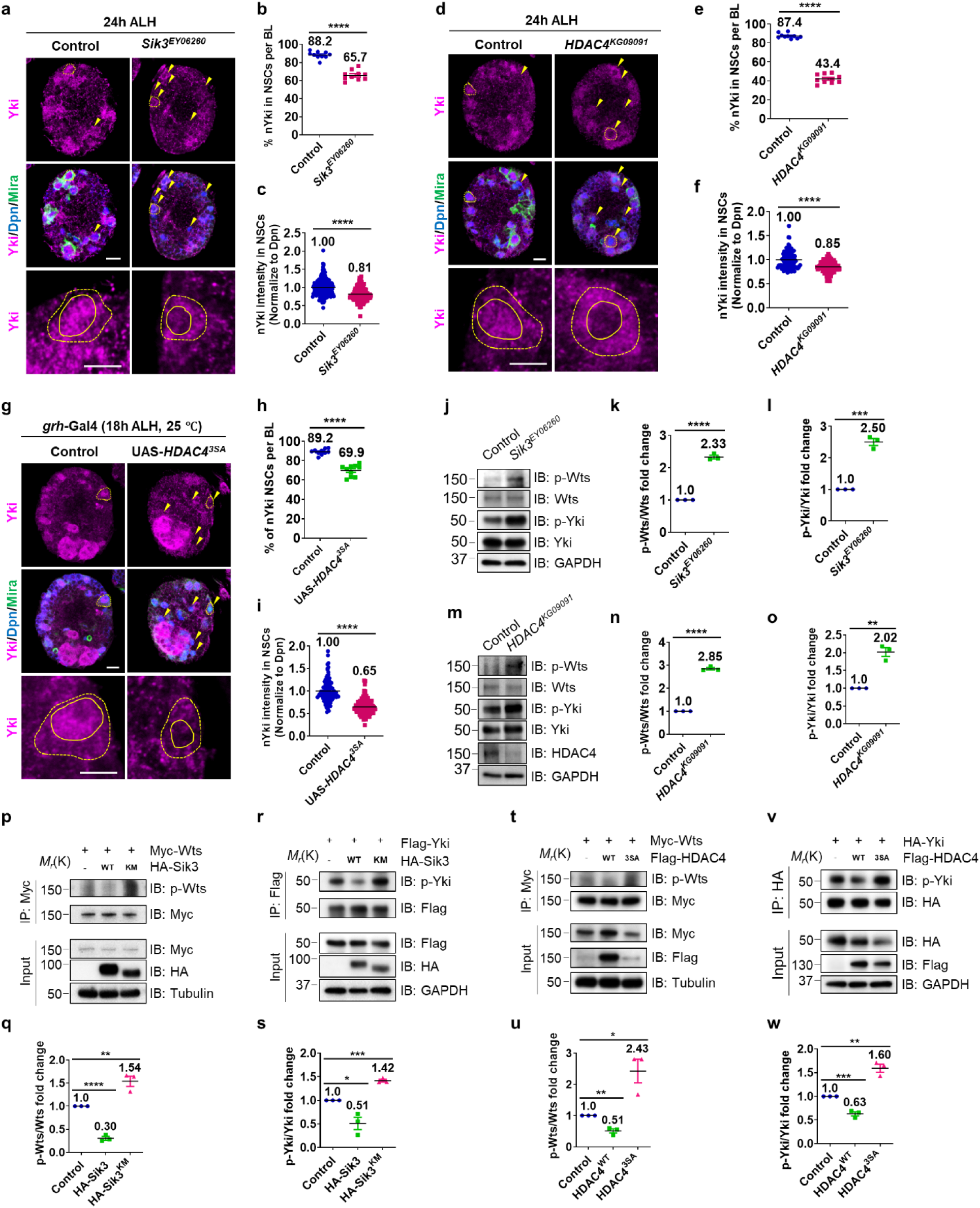
SIK3-HDAC4 promotes Yki nuclear localization by suppressing Wts phosphorylation and kinase activit. **a, d, g** At 24 h or 18 h ALH, larval brain lobes were labeled with indicated proteins. NSCs were marked by Dpn and Mira. Yellow arrows point to NSCs without nYki localization. Yellow dotted circles label the NSCs, yellow circles label the nucleus of the NSCs. Lower panels are enlarged views of the NSCs marked by yellow dotted circles in the upper panels, the. **b, e, h** Quantification of NSCs with nYki per BL in (**a, d, g**) respectively. **b** Control (yw), 88.2 ± 3.9, n = 10; *Sik3^EY06260^*, 65.7 ± 5.4, P = 3.3E−09, n = 10. **e** Control (yw), 87.4 ± 2.7, n = 10; *HDAC4^KG09091^*, 43.4 ± 4.6, P = 5.5E−16, n = 10. **h** Control (β-galRNAi), 89.2 ± 2.9, n = 10; UAS-HDAC43SA, 69.9 ± 5.9, P = 3.0E-8, n = 10. **c, f, i** Quantification of nYki intensity (normalized to Dpn) in NSCs in (**a, d, g**) respectively. (**c**) Control, 1 ± 0.24, n = 10; *Sik3^EY06260^*, 0.81 ± 0.18, P = 1.3E-14, n = 10. **f** Control, 1 ± 0.18, n = 10; *HDAC4^KG09091^*, 0.85 ± 0.12, P = 3.2E-11, n = 10. **i** Control, 1 ± 0.25, n = 10; UAS-HDAC43SA, 0.65 ± 0.18, P = 1.3E-23, n = 10. **j, m** At 24 h ALH, the larval brains from *Sik3^EY06260^* and *HDAC4^KG09091^* were dissected, proteins extracted from the brains were used for Western blots. **k, n** Quantification of the fold-change of p-Wts/Wts, n=3. **k** Control (yw), 1; *Sik3^EY06260^*, 2.33 ± 0.14, P = 1.8E-05. **n** Control (yw), 1; *HDAC4^KG09091^*, 2.85 ± 0.09, P = 3.8E-06. **l, o**Quantification of the fold-change of p-Yki/Yki, n=3. **l** Control (yw), 1; *Sik3^EY06260^*, 2.5 ± 0.19, P = 0.0002. **o** Control (yw), 1; *HDAC4^KG09091^*, 2.02 ± 0.21, P = 0.0011. **p, r, t, v** S2 cells transfected with indicated constructs were harvested for IP and Western blot. **q, s, u, w** Quantifications for (**p, r, t, v**), respectively. **q** Control, 1; *Sik3*, 0.3 ± 0.07, P = 6.2E-05; *Sik3KM*, 1.54 ± 0.19, P = 0.009. **s** Control, 1; *Sik3*, 0.51 ± 0.22, P = 0.02; *Sik3KM*, 1.42 ± 0.05, P = 0.0001. **u** Control, 1; HDAC4, 0.51 ± 0.12, P = 0.002; HDAC43SA, 2.43 ± 0.65, P = 0.02. **w** Control, 1; HDAC4, 0.63 ± 0.07, P = 0.0009; HDAC43SA, 1.6 ± 0.14, P = 0.002. Data are presented as mean ± SD. * for 0.05≤ P ≤ 0.01, ** for P ≤ 0.01, *** for P ≤ 0.001, **** for P ≤ 0.0001. Scale bars: 10 μm.

### SIK3 and HDAC4 inhibit Wts phosphorylation and kinase activity

Phosphorylation of Wts at Thr1077 (p-Wts) by the Hpo kinase is regarded as a principal readout of Hippo pathway activity ^98^. Upon phosphorylation by Hpo, Wts in turn phosphorylates Yki primarily at Ser168 (p-Yki), resulting in Yki cytoplasmic retention and functional inactivation ^35,98,99^. We therefore examined p-Wts and p-Yki levels *in vivo* in *Drosophila* larval brains. At 24 h ALH, larval brains from *Sik3* and *HDAC4* mutants exhibited a significant increase in p-Wts levels by 2.33-fold and 2.85-fold respectively, compared with controls (Figure 6j–k, m–n). Consistently, p-Yki levels were also significantly elevated in *Sik3* and *HDAC4* mutant brains by 2.50-fold and 2.02-fold respectively (Figure 6j, l and m, o). These *in vivo* findings were corroborated by *in vitro* experiments in S2 cells, in which dsRNA-mediated depletion of *Sik3* or *HDAC4* significantly increased the phosphorylation levels of both Wts and Yki (Supplementary Figure 5a–f). In contrast, overexpression of wild-type *Sik3* or *HDAC4* in S2 cells led to a significant reduction in p-Wts and p-Yki levels, whereas overexpression of kinase-dead SIK3 (*Sik3^K70M^*) or SIK3 phosphorylation–defective HDAC4 (*HDAC4^3SA^*) resulted in a marked increase in p-Wts and p-Yki (Figure 6p–w). However, overexpression of either wild-type or SIK3 phosphorylation–defective HDAC4 did not affect Hpo phosphorylation at T195 by Tao-1, which is required for Hpo kinase activity (Supplementary Figure 5g, h). Collectively, these data indicate that the SIK3–HDAC4 axis suppresses Wts phosphorylation and kinase activity, thereby promoting Yki nuclear accumulation and NSC reactivation.

### SIK3-HDAC4 promotes the expression of Hippo pathway target genes in NSCs

Next, we assessed the expression of known Yki targets during NSC reactivation, including cycE and bantam, as readouts of Yki activity. Consistent with reduced Yki function, loss of *Sik3* or *HDAC4* in larval brains at 24 h ALH resulted in a marked decrease in CycE intensity in NSCs (Supplementary Figure 5i–l). In contrast, at 24 h ALH, overexpression of wild-type *HDAC4* in NSCs significantly increased *bantam*-lacZ reporter activity, whereas overexpression of the *Sik3* phosphorylation–defective mutant *HDAC4^3SA^*significantly reduced *bantam*-lacZ expression (Supplementary Figure 6a, b). Furthermore, EdU incorporation assays revealed that overexpression of *bantam* significantly rescued the NSC reactivation defects induced by *HDAC4^3SA^*overexpression (Supplementary Figure 6c, d). Together, these findings provide functional evidence that the SIK3–HDAC4 axis inhibits Hippo signaling, enabling Yki-dependent gene expression during NSC reactivation.

### HDAC4 promotes NSC reactivation by inhibiting the Hippo pathway

To establish epistasis between HDAC4 and the Hippo pathway in NSC reactivation, we performed genetic analyses. At 24 h ALH, knockdown of *wts* significantly suppressed reactivation phenotypes observed in *HDAC4*-RNAi brains shown by EdU incorporation, reducing the proportion of EdU-negative quiescent NSCs from 29.2% in *HDAC4*-RNAi alone to 11.8% in *HDAC4*; *wts* double knockdown brains (Figure 7a, b). *wts*-RNAi alone was undistinguishable from the control, with the percentage of EdU-negative NSCs comparable to controls (7.5% vs. 7.9%; Figure 7a, b). Similarly, overexpression of a constitutively active Yki mutant (Yki^S168A^) markedly alleviated the NSC reactivation defects caused by HDAC4 knockdown, decreasing the percentage of EdU-negative quiescent NSCs from 29.2% to 11.3% (Figure 7a, b). Moreover, both *wts* knockdown and *yki^S168A^* overexpression significantly suppressed the NSC reactivation defects induced by HDAC4^K902R^, which appeared to act as a dominant-negative form. Specifically, the percentage of EdU-negative quiescent NSCs decreased from 34.2% in UAS-*HDAC4^K902R^* to 20.6% with *wts*-RNAi and to 19.5% with *yki^S168A^* overexpression (Figure 7c, d). These results strongly support the conclusion that HDAC4 promotes NSC reactivation by inhibiting Wts activity.

**Figure 7.**
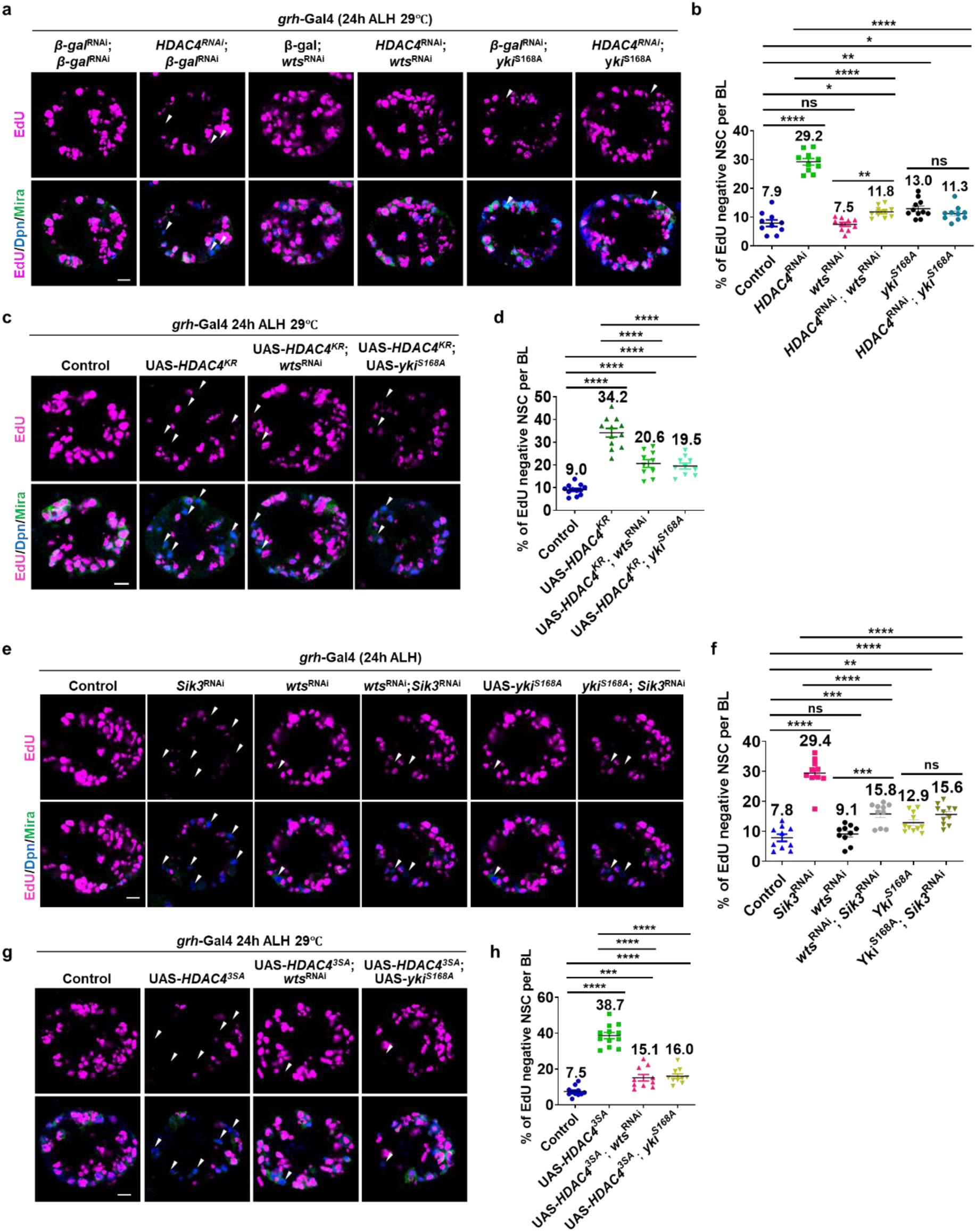
SIK3-HDAC4 promotes NSC reactivation by inhibiting the Hippo pathway a, c, e, **g** At 24 h ALH, larval brain lobes of indicated genotypes were analyzed for EdU incorporation. NSCs were marked by Dpn and Mira. White arrows point to EdU negative quiescent NSCs. **b, d, f, h** Quantification graph of EdU- NSCs per BL for genotypes in (**a, c, e, g**) respectively. **b** Control (β-galRNAi; β-galRNAi), 7.9 ± 3.6, n = 10; *HDAC4*RNAi; β-galRNAi, 29.2 ± 3.6, P = 1.0E-10, n = 10; β-galRNAi; *wts*RNAi, 7.5 ± 2.2, P = 0.77, n = 10; *HDAC4*RNAi; *wts*RNAi, 11.8 ± 2.0, P = 0.008, P = 2.3E-07, P = 0.002, n = 10; β-galRNAi; *ykiS168A*, 13 ± 3.3, P = 0.006, n = 11; *HDAC4*RNAi; *ykiS168A*, 11.3 ± 2.6, P = 0.06, P = 1.6E-10, P = 0.2, n = 10. **d**Control (β-galRNAi; β-galRNAi), 9 ± 2, n = 12; UAS-*HDAC4KR*; β-galRNAi, 34.2 ± 6.8, P = 2.1E-11, n = 12; UAS-*HDAC4KR*; *wts*RNAi, 20.6 ± 5.4, P = 1.6E-06, P = 4.5E-05, n = 10; UAS-*HDAC4KR*; *ykiS168A*, 19.5 ± 4.1, P = 3E-07, P = 6E-06, n = 10. **f** Control (β-galRNAi; β-galRNAi), 7.8 ± 3.8, n = 10; β-galRNAi; *Sik3*RNAi; 29.4 ± 5.1, P = 2.8E-09, n = 10; *wts*RNAi; β-galRNAi, 9.1 ± 3.1, P = 0.4, n = 10; *wts*RNAi; *Sik3*RNAi, 15.8 ± 3.8, P = 0.0002, P = 0.0004, P = 2.3E-06, n = 10; *ykiS168*A; β-galRNAi, 12.9 ± 3.4, P = 0.004, n = 11; *ykiS168A*; *Sik3*RNAi, 15.6 ± 3.4, P = 6.6E-05, P = 4.7E-07, P = 0.06, n = 11. **h** Control (β-galRNAi; β-galRNAi), 7.5± 2.9, n = 11; UAS-*HDAC43SA*; β-galRNAi, 38.7 ± 6.7, P = 5.6E-13, n = 12; UAS-*HDAC43SA*; *wts*RNAi, 15.1 ± 5.8, P = 0.0009, P = 1.3E-08, n = 10; UAS-*HDAC43SA*; *ykiS168*A, 16 ± 4.2, P = 2.3E-05, P = 3.5E-09, n = 10. Data are presented as mean ± SD. ns for P > 0.05, * for 0.05≤ P ≤ 0.01, ** for P ≤ 0.01, *** for P ≤ 0.001, **** for P ≤ 0.0001. Scale bars: 10 μm.

### SIK3 promotes NSC reactivation by inhibiting the Hippo pathway

Finally, we performed epistasis analyses between SIK3 and the Hippo pathway in NSC reactivation. At 24 h ALH *wts*-RNAi significantly suppressed the reactivation phenotype observed in *Sik3*-RNAi brains, reducing the proportion of quiescent NSCs from 29.4% in *Sik3*-RNAi alone to 15.8% in *wts*; *Sik3* double knockdown brains (Figure 7e, f). Similarly, overexpression of *yki^S168A^*significantly alleviated the NSC reactivation defects caused by *Sik3* depletion, decreasing the percentage of EdU-negative NSCs from 29.4% to 15.6% (Figure 7e, f). Likewise, NSC reactivation defects caused by overexpression of the SIK3 phosphorylation–defective mutant *HDAC4^3SA^* were significantly suppressed by either *wts*-RNAi or *yki^S168A^* overexpression (Figure 7g, h; UAS-*HDAC4^3SA^*, 38.7%; UAS-*HDAC4^3SA^*; *wts*-RNAi, 15.1%; UAS-*HDAC4^3SA^*; UAS-*yki^S168A^*, 16.0%). Together, these genetic interaction analyses provide strong evidence that the SIK3–HDAC4 axis promotes NSC reactivation by inhibiting Hippo pathway signaling (Figure 8).

**Figure 8.**
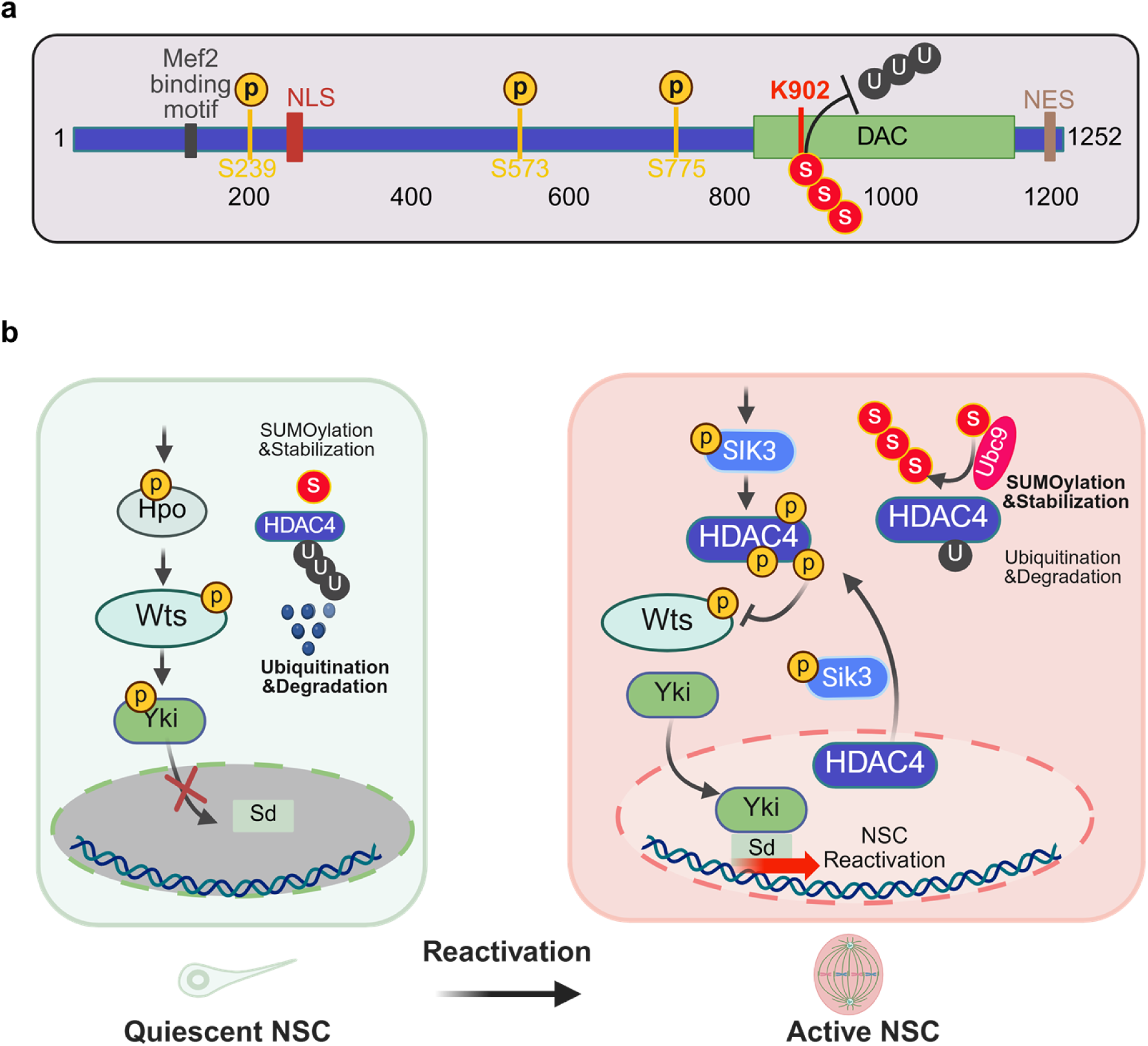
Working model of SIK3-HDAC4-Wts in promoting quiescent NSC reactivation. **a** Diagrammatic illustration of HDAC4 protein domains and PTM sites. MEF2: Mef2 binding domain; NLS: nuclear localization signal; phosphorylation sites by SIK3: S239: Serine 239 residue; S573: Serine 573 residue; S775: Serine 775 residue; DAC: deacetylase domain; K902: Lysine 902 residue, HDAC4 SUMOylation site; S: SUMO; U: Ubiquitin; NES: nuclear export signal. **b** In quiescent NSCs, the main kinase of the Hippo pathway Wts is activated and phosphorylates Yki to sequester Yki in the cytoplasm to maintain NSC quiescence. Basal levels of HDAC4 could be maintained by ubiquitination marks for degradation. During NSC reactivation, SIK3 and HDAC4 expression levels increase alongside the activation of the SUMO pathway, which promotes HDAC4 SUMOylation and stabilization by inhibiting its ubiquitination and proteasomal degradation. SIK3 directly interacts with and phosphorylates HDAC4, facilitating its cytoplasmic retention and enabling efficient binding to Wts. This interaction suppresses Wts phosphorylation and kinase activity, leading to enhanced nuclear translocation of Yki and ultimately triggering the reactivation of quiescent NSCs.

## Discussion

In this study, we identify a previously unrecognized SIK3–HDAC4–Warts (Wts) signaling axis that functions as a molecular gatekeeper controlling NSC reactivation in *Drosophila*. Our findings uncover HDAC4 as a central integrator of kinase signaling that suppresses the Hippo pathway activity to enable Yki-dependent transcription and cell cycle re-entry. SUMOylation of HDAC4 leads to its protein stabilization by preventing it from ubiquitination. SIK3 phosphorylates HDAC4, facilitating its cytoplasmic retention and enabling its efficient association with Wts. HDAC4 interaction with Wts prevents Wts phosphorylation and consequently suppresses its kinase activity. This leads to enhanced nuclear accumulation of Yki to ultimately trigger the reactivation of quiescent NSCs. Genetic epistasis analyses place SIK3 and HDAC4 upstream of Wts and Yki, establishing the SIK3–HDAC4-Wts-Yki axis in NSCs.

We show that HDAC4 loss-of-function results in a microcephaly-like phenotype in *Drosophila*. In contrast, a significant upregulation of hHDAC4 in glioma patients was found to promote glioma cell proliferation and invasion ^100^. Our data reveals that HDAC4 activity is tightly regulated at multiple levels, including protein abundance, post-translational modification and subcellular localization. Our previous work demonstrated that Wts SUMOylation results in the inactivation of Wts for NSC reactivation ^39^. In this study, we show that HDAC4, a novel regulator of NSC reactivation, is also SUMOylated. We find that HDAC4 protein levels increase during NSC reactivation, partially contributed by SUMOylation at HDAC4-Lys902, which stabilizes HDAC4 by antagonizing ubiquitin-mediated proteasomal degradation. Surprisingly, Lys902 functions as both a major SUMOylation site and a critical ubiquitination site on HDAC4. When HDAC4 is SUMOylated, its ubiquitination is precluded, resulting in HDAC4 stabilization. Conversely, in the absence of SUMOylation, ubiquitination at this residue promotes its proteasomal degradation. Therefore, Lys902 likely functions as a competitive modification site for SUMO and ubiquitin, serving as a molecular switch that regulates HDAC4 protein stability. Low levels of HDAC4 in quiescent NSCs is likely insufficient to inhibit the Hippo pathway, thus maintaining NSC quiescence. As expression levels of HDAC4 and SUMO increase, HDAC4 levels are further augmented by its SUMOylation, crossing the threshold of HDAC4 protein levels required for the inhibition of the Hippo pathway. This SUMOylation–ubiquitination antagonism represents a rapid and reversible mechanism by which NSCs fine-tune the timing and robustness of their reactivation response. However, we do note that human HDAC4 was reported to be SUMOylated at Lys-559 ^72^, a site is absent in *Drosophila* HDAC4. Intriguingly, we found that human HDAC4-K737 residue, the equivalent of *Drosophila* HDAC4-K902, was predicted to be a ubiquitylation site instead of SUMOylation site (Supplementary Table 4, 5). Further study is required to determine if human HDAC4 is ubiquitinated at this site to moderate its stability like the *Drosophila* system.

In addition to protein stability, the subcellular localization of HDAC4 in NSCs emerges as a critical determinant of function. The HDAC4^3SA^ mutant in both vertebrates and *Drosophila* results in nuclear retention of HDAC4 ^52,55,101,102^, which was found to be strongly associated with Alzheimer’s disease and Parkinson’s disease ^103–105^. SIK3, a ubiquitously expressed serine/threonine kinase belonging to the AMPK family, regulates diverse cellular processes including metabolism, innate immunity, bone formation, and hippocampal synaptic plasticity ^106–108^. Dysregulation of SIK3 has been identified in various cancer types and Alzheimer’s disease (AD) ^95,109^. We show that SIK3-mediated phosphorylation promotes HDAC4 cytoplasmic retention, which is essential for its interaction with Wts. In contrast, phosphorylation-defective HDAC4 (HDAC4^3SA^) accumulates in the nucleus, fails to associate efficiently with Wts, and thus severely impairs NSC reactivation. Given our findings on *Drosophila* SIK3 in NSC reactivation, it will be of great interest to investigate whether mammalian SIK3 could regulate adult neurogenesis, a process that is often impaired in mouse models of AD.

In this study, we showed that HDAC4 physically associates with C-terminal region of Wts and prevents its activation. Since this C-terminal half of Wts contains a phosphorylation site (T1077) critical for its activation by Hpo ^110^,we speculate that the HDAC4-Wts association likely sterically hinders Wts phosphorylation by Hpo, thereby preventing its activation. Alternatively, since the kinase domain of Wts is also located within its C-terminal region, the HDAC4-Wts association might also interfere with the potential interaction between Wts kinase and its substrate. HDAC4 is a class IIa histone deacetylase that is known to exhibit intrinsically weak deacetylase activity toward canonical histone substrates, attributed to a conserved tyrosine-to-histidine substitution in its catalytic domain ^111^. Instead, HDAC4 primarily functions as a signal-responsive transcriptional repressor. In this study, we demonstrated that HDAC4 inhibits the Hippo pathway to promote NSC reactivation via an association with Wts, a novel mechanism that is distinct from its canonical deacetylation or transcriptional repressor function. The nucleocytoplasmic shuttling of HDAC4 is governed by SIK3/CaMK phosphorylation-dependent 14-3-3 binding, which is known to promote the cytoplasmic sequestration of HDAC4 in mammalian fibroblasts and HEK293 cells^90^. Additionally, HDAC4 deacetylates non-histone targets such as MEF2, thereby modulating their transcriptional activity independently of its canonical deacetylase function ^112^. However, we observed how increased HDAC4 nuclear localization in HDAC4^3SA^ mutants resulted in defects in NSC reactivation, suggesting that the transcriptional repressor function of HDAC4 might be dispensable for NSC reactivation. It remains to be tested whether HDAC4 deacetylase activity is required for NSC reactivation.

The components of this SIK3-HDAC4-Wts axis are highly conserved across evolution, underscoring the broader relevance of our findings. SIK3, HDAC4, and the core Hippo pathway are conserved from flies to humans, and analysis of single-cell RNA-sequencing datasets reveals that both SIK3 and HDAC4 are enriched in the NSCs of both the fly and human fetal brain. In mammals, HDAC4 dysregulation is associated with a wide spectrum of neurological disorders, including intellectual disability ^41,55^, autism spectrum disorders ^59,63^, Alzheimer’s disease ^50,59–62^, Huntington’s disease ^64^, and Parkinson’s disease ^103^. Likewise, YAP/TAZ—the mammalian homologs of Yki play well-established roles in neural stem cell activation ^38^, gliogenesis ^113^, and tumorigenesis ^114^. There is also emerging evidence for the implication of the Hippo pathway in neurodegenerative diseases^115^. Therefore, there is a strong indication that the dysfunction of the HDAC4-Wts axis might similarly contribute to these neurodevelopmental and neurodegenerative diseases.

In summary, we propose a model in which NSC reactivation is driven by coordinated upregulation and stabilization of HDAC4 through SIK3-dependent phosphorylation and cytoplasmic retention of HDAC4. Wts activity is thus consequently inhibited. This cascade culminates in the nuclear accumulation of Yki and subsequent transcriptional activation of proliferative programs. This discovery of the SIK3-HDAC4-Wts axis advances our understanding of stem cell state transitions and provides new insight into conserved mechanisms underlying brain development and neurological disease.

## Materials and Methods

### Fly stocks and genetics

Fly stocks and genetic crosses were raised at 25°C unless otherwise stated. Fly stocks were fed with standard fly food (0.8% *Drosophila* agar, 5.8% Cornmeal, 5.1% Dextrose, and 2.4% Brewer’s yeast). The following fly stocks were used in this study: UAS-Flag-HDAC4 (Biao Wang), pUAST-Flag-HDAC4^3SA^ (Biao Wang), *grh*-Gal4 (Andrea Brand), *tub*-Gal4, pUAST-Flag-HDAC4^RNAi^ Resistant (generated in this work by WellGenetics). UAS-Venus HDAC4^K902R^ (generated in this work), UAS-Myc-Sik3 (Jongkyeong Chung), UAS-Myc-Sik3^K70M^ (Jongkyeong Chung). The following fly strains were obtained from BDSC: *HDAC4^KG09091^*(15159), *HDAC4^EY06270^* (16708), UAS-*InR* (8440), *smt3^04493^*(11378), *β*-*gal*^RNAi^ (50680), UAS-CD8-GFP (#32186), *lwr^13^* (9323), *wts*^RNAi^ (34064), UAS-*yki^S168A^* (28818), Ban-lacZ (10154), UAS-*bantam* (60671), *Sik3*^RNAi^- 1(28366), *Sik3^EY06260^*(15962) and *Sik3^f03280^* (18634). RNAi lines including *HDAC4*^RNAi^-1 (330055), *HDAC4*^RNAi^-2 (20522), *smt3^RNAi^* (105980), *Sik3*^RNAi^-2 (39866), and *Sik3*^RNAi^-3 (106268) were obtained from the Vienna *Drosophila* Resource Center (VDRC). UAS-*β*-*gal*^RNAi^ is often used as a control UAS element to balance the total number of UAS elements in each genotype. Various RNAi knockdown or overexpression constructs were induced using *grh*-Gal4 unless otherwise stated. All experiments were carried out at 25°C, except for RNAi knockdown or overexpression experiments that were performed at 29°C, unless otherwise indicated.

### Plasmids and transgenic flies

pAc-Myc-Smt3 and pAc-Myc-Smt3^AA^ ^39^, pUAST-Flag-HDAC4 and pUAST-Flag-HDAC4^3SA^ (Helen Fitzsimons), pAc-Myc-Wts ^40^, pAc-Myc-Wts^NT^ (1-603aa), pAc-Myc-Wts^CT^ (601-1105aa), pUAST-HA-Sik3 and pUAST-HA-Sik3^K70M^ (Addgene).

Point mutation plasmids were generated using wild-type plasmids as templates, and the PerfectStart® Taq DNA Polymerase kit (TransGen Biotech) was used to perform PCR. The following primers were used for generating point mutation plasmids:

HDAC4^K902R^ F: TGCCAGCTCAGCAGGCCCAGGTTGGAAAACAC

HDAC4^K902R^ R: CTGGGCCTGCTGAGCTGGCACTGATTCGAAC

Double strand RNA primers for gene knockdown are as below:

HDAC4-dsRNA-1 F: TAATACGACTCACTATAGGG<u>TTGCAGTGCGCAACAATAAT</u>

HDAC4-dsRNA-1 R: TAATACGACTCACTATAGGG<u>TATGCAAGCCACGTAATTGC</u>

HDAC4-dsRNA-2 F: TAATACGACTCACTATAGGG<u>GCCAGCACCGCCAGCTAATG</u>

HDAC4-dsRNA-2 R:TAATACGACTCACTATAGGG<u>CCGCACGACAAACGCACAAA</u>

Sik3-dsRNA F-1: TAATACGACTCACTATAGGG <u>GCGAGCAGCTAATACGGAAC</u>

Sik3-dsRNA R-1: TAATACGACTCACTATAGGG <u>TATTTCTCCAGATCCACGGC</u>

Sik3-dsRNA F-2: TAATACGACTCACTATAGGG <u>TCAATGGTGCGAATACCAGA</u>

Sik3-dsRNA R-2: TAATACGACTCACTATAGGG <u>GTCAACCTTTGCAGCTCCTC</u>

pUAST-Flag-HDAC4^RNAi^ resistant fly line was designed to resistant to VDRC#20522 was generated by WellGenetics. Transgenic flies: UAS-*venus*-*HDAC4^K902R^* lines were generated by P-element-mediated transformation (BestGene).

### Quantitative real-time PCR

Total RNA was isolated from *Drosophila* larva brain with TRI reagent (Sigma Aldrich) followed by DNase I treatment (Sigma-Aldrich) as described by the manufacturer. The quantitative (q)PCR was performed using Maxima SYBR Grebb/ROX qPCR Master Mix (Fermentas) according to the manufacturer’s protocol. Primers used here are as below:

Smt3-qF: AGAAGGGAGGTGAGACCGAG

Smt3-qR: GAGTGTCGTTCTCGTTGATG

HDAC4 qF: GAACTCCTTCAGTTGGCCAAC

HDAC4 qR: GGTTTCATCCTGATCCATCG

### EdU (5-ethynyl-20-deoxyuridine) incorporation assay

Larvae were fed with food containing 2% (0.2 mM) EdU from Click-iT EdU Imaging Kits (Invitrogen) for 4 h before dissection. The dissected larval brains were fixed with 4% EM grade formaldehyde for 22 min, followed by washing thrice with 0.3% PBST (PBS + 0.3% Triton-100) and blocking with 3% BSA (bovine serum albumin) in PBST for 45 min. Following blocking, incorporated EdU was detected by Alexa Fluor azide, according to the Click-iT EdU protocol (Invitrogen). The brains were then washed briefly twice, blocked with 3% BSA again for 20 min, and subjected to immunohistochemistry.

### Immunohistochemistry

*Drosophila* larvae were dissected in phosphate-buffered saline (PBS), and the larval brains were fixed in 4% EM-grade formaldehyde in PBST for 22 min. After washing thrice with 0.3% PBT (10 min each), brain samples were blocked with 3% BSA in 0.3% PBST for 45 min. Blocked brain samples were incubated with primary antibodies diluted in 3% BSA overnight at 4°C. Following this, they were again washed thrice with 0.3% PBST (10 min each) and incubated with secondary antibodies diluted in 0.3% PBT for 1.5h. DNA was labeled with DAPI (1:1,500, Molecular Probes), and after washing twice with 0.3% PBT (10 min each), larval brains were mounted onto Vector shield (Vector Laboratory) for Confocal microscopy. Confocal images were taken by a Zeiss LSM710 confocal microscope and processed with ImageJ for brightness and contrast adjustment. Primary antibodies used were as follows: guinea pig anti-Dpn (1:1,000, J Skeath), mouse anti-Mira (1:50, F. Matsuzaki), rabbit anti-PH3 (1:500, Cell signaling), rabbit anti-HDAC4 (1:100, generated in this work by Abmart), mouse anti-GFP (1:2,000, Fengwei Yu), rabbit anti-Yki (1:100, generated by GenScript), mouse anti-β-gal (1:100, Promega) and mouse anti-CycE (1:200, H. Richardson), anti-Myc antibody (1:2,000, abcam) and rabbit anti-HA (1:2,000, Sigma-Aldrich). The secondary antibodies used were conjugated with Alexa Fluor 488, 555 or 647 (Jackson laboratory).

### Cell lines, transfection and drug treatment

*Drosophila* S2 cells (CVCL_Z232), originally from William Chia’s laboratory, were cultured in Express Five serum-free medium (Gibco) supplemented with 2 mM Glutamine (Thermo Fisher Scientific). For transient expression of proteins, S2 cells were transfected using Effectene Transfection Reagent (QIAGEN) according to the manufacturer’s protocol. For RNAi experiments, *Drosophila* S2 cells were cultured and incubated with dsRNA for 72 h. For inhibition of SUMOylation, S2 cells were treated with DMSO as control and the ML-792 (0.2 µM) inhibitor as experiment group for 6h before harvest. For cycloheximide (CHX) chase assays, S2 cells were treated with CHX at 2 h intervals, with the longest drug treatment being for 6 h.

### Protein extraction, immunoprecipitation, immunoblotting and immunofluorescence

S2 cells were transfected with the indicated plasmids and cultured for 48 h; alternately *Drosophila* larval brains were dissected at certain time points, and were collected and lysed in NP-40 lysis buffer along with protease inhibitors (Roche) for 30 min on a rotor at 4 °C. Immunoprecipitation was performed using mouse anti-Flag antibody (1:200, Sigma), rabbit anti-HDAC4 (1:100, generated in this work by Abmart), mouse anti-Myc antibody (1:200, abcam) or rat anti-HA (1:100, Roche) and protein G Sepharose beads according to manufacturer’s instruction. The samples were separated by SDS–PAGE and analyzed by Western blotting. Blots were probed with the following antibodies: rabbit anti-HDAC4 (1:500, generated in this work by Abmart), rabbit anti-GAPDH1 (1:500, GeneTex), rabbit anti-SUMO (1:500, Albert J. Courey), mouse anti-Flag antibody (1:2,000, Sigma), mouse anti-Myc antibody (1:2,000, abcam), mouse anti-Tubulin (1:2,000, Sigma-Aldrich), rabbit anti-p-HDAC4 (Ser239, 1:500, Cell Signaling Technology), rabbit anti-Wts (1:500, generated in this work by Abmart), rabbit anti-HA (1:2,000, Sigma-Aldrich), rabbit anti-p-Wts (1:500, Duojia Pan), rabbit anti-Wts (1:500, Abmart), rabbit anti-p-Yki (1:2,000, Duojia Pan), rabbit anti-Yki (1:2,000, generated by GenScript) and p-Hpo (p-Mst1, 1:1,000, Cell Signaling Technology). The secondary antibodies used were conjugated with HRP (Horseradish Peroxidase). For immunofluorescence (IF) assay, cells were transfected with the indicated plasmids after being seeded on coverslips. 48h later, cells were washed with PBS and fixed in 4% paraformaldehyde for 20 min, and permeabilized with 0.5% Triton X-100 for 30 min at room temperature. Cells were then blocked for 30 min in 3% BSA and incubated with primary antibodies for 2 h, and then with fluorophore-conjugated secondary antibodies for another 1 h at room temperature. They were then treated with DAPI (1:1,500, Molecular Probes) for 6 min to mark nuclear DNA before the slides were mounted. Cell images were captured with Zeiss LSM710 Confocal microscope. Antibodies used for IF: rabbit anti-HDAC4 (1:1000, generated in this work by Abmart), Phalloidin (1:1,000, Invitrogen) mouse anti-Flag antibody (1:2,000, Sigma), rabbit anti-HA (1:1,000, Sigma-Aldrich), mouse anti-Myc antibody (1:2,000, abcam). The secondary antibody used was conjugated with Alexa Fluor 488 (Jackson laboratory).

### Proximity ligation assay (PLA)

Proximity ligation assay (PLA) is based on the principle that secondary antibodies conjugated to PLA PLUS or PLA MINUS probes bind to anti-Flag and anti-Myc primary antibodies, respectively. During the ligation step, connector oligonucleotides hybridize to the PLA probes, and T4 DNA ligase catalyzes their joining to generate a circular DNA template. This circularized template is subsequently amplified by DNA polymerase, and the amplification products are detected by fluorescently labeled complementary oligonucleotides, enabling protein–protein interactions to be visualized as discrete PLA puncta within cells (adapted from Duolink PLA, Merck). PLA was performed in S2 cells transfected using Effectene Transfection Reagent (QIAGEN) with the following plasmids: control Myc, control Flag, Myc-Wts, Flag-HDAC4, Flag-HDAC4^K902R^, and Flag-HDAC4^3SA^. Cells were washed three times with cold PBS, fixed in 4% EM-grade formaldehyde in PBS for 15 min, and blocked with 5% BSA in PBS-T (0.1% Triton X-100) for 45 min. Primary antibody incubation was carried out at room temperature for 2 h, after which the Duolink PLA procedure (Sigma-Aldrich) was performed according to the manufacturer’s instructions. Following primary antibody incubation, cells were incubated with PLA probes at 37 °C for 1 h, washed twice with Buffer A for 5 min each at room temperature, and subjected to probe ligation at 37 °C for 30 min. Signal amplification was performed at 37 °C for 100 min, followed by two washes with Buffer B for 10 min each at room temperature. Cells were then washed once with 0.01× Buffer B and incubated with primary antibodies diluted in 3% BSA in PBS for 2 h at room temperature. After two washes with 0.1% PBS-T, cells were incubated with secondary antibodies for 1.5 h at room temperature and finally mounted using in situ mounting medium containing DAPI (Duolink, Sigma-Aldrich).

### SUMOylation assay and ubiquitination assay

For SUMOylation assays, the lysis buffer was supplemented with the protease inhibitor N-ethylmaleimide (NEM; 10 mM final concentration; Sigma) to suppress the deconjugating activity of SUMO proteases. Protein extracts from S2 cells or larval brains were prepared as described above in the presence of the SUMO isopeptidase inhibitor NEM, followed by cell lysis and subsequent immunoprecipitation and Western blot analysis as previously described. For ubiquitination assay, S2 cells were treated with 10 µM MG132 for 4 h before harvest. Then followed by cell lysis and subsequent immunoprecipitation and Western blot analysis as previously described.

### Generation of Rabbit anti-HDAC4 antibodies

The N-terminal coding sequence (encoding 1–300 amino acids) of HDAC4 was amplified by PCR and inserted into a GST-containing vector. The purified GST-HDAC4^1-300^ was then injected into rabbits. Rabbit anti-HDAC4 antibodies were generated and purified by Abmart (Shanghai).

### Single cell RNA-seq data analysis

Raw data was downloaded from GSE134722 Brunet et. al. ^67^ and processed by Seurat 4.0. The raw data was first analyzed according to the methods in Brunet et. al. ^67^. Sub-clustering was then performed on the neural stem cells cluster. Quiescent and active neural stem cells were annotated by the expression of proliferating markers: Wor, CycA, CycE and PCNA. Human fetal brain scRNA-seq dataset with cell-type annotations from human fetal brain during peak neurogenesis and early gliogenesis ^69^ was re-analyzed. Seurat v5 and R 4.4.0 were used to examine gene expression level among all the neuronal cells and glial cells. SOX2, an NSC marker; PAX6, an apical radial glia cells marker; HOPX, a basal radial glia cells marker, were included in the analysis as a control.

### Quantification and statistical analysis

Statistical analysis was performed using GraphPad Prism 6. *P*-values were calculated by performing two-tailed, unpaired Students’ T-test. In all graphs, ^∗^indicates 0.05 < p < 0.01, ^∗∗^indicates p < 0.01, ^∗∗∗^indicates p < 0.001, ^∗∗∗∗^indicates p < 0.0001, ns indicates p > 0.05. Cell population numbers were quantified and represented as mean ± SD and/or percentage of phenotype. Each experiment has at least three independent replications. We distinguish active NSCs from quiescent NSCs based on the cell size, as active NSCs are larger than 7 µm.

## Acknowledgments

We thank Duojia Pan, Fengwei Yu, Shian Wu, Biao Wang, Jongkyeong Chung, Helen Fitzsimons, Albert J. Courey and the Bloomington *Drosophila* Stock Center, Vienna *Drosophila* Resource Center, Kyoto Stock Centre DGGR, and the Developmental Studies Hybridoma Bank, Abmart INC for plasmids, fly stocks and antibodies. This work is supported by the Ministry of Health-Singapore National Medical Research Council YIRG MOH-000924 to Yang Gao and Duke-NUS Khoo Postdoctoral Fellowship Award KPFA/2025/0080 to Amanda Ng.

## Author contributions

Conceptualization: Y.G. and H.W Methodology: Y.G., J.L. and A.N.Y.E. Investigation: Y.G.

Visualization: Y.G. Supervision: H.W. Writing—original draft: Y.G.

Writing—review & editing: Y.G., A.N.Y.E. and H.W.

## Data and materials availability

The authors declare that the data supporting the findings of this study are available within the paper and the source data for figures are provided with the paper.

## Figures and Tables

**Supplementary Figure 1.**
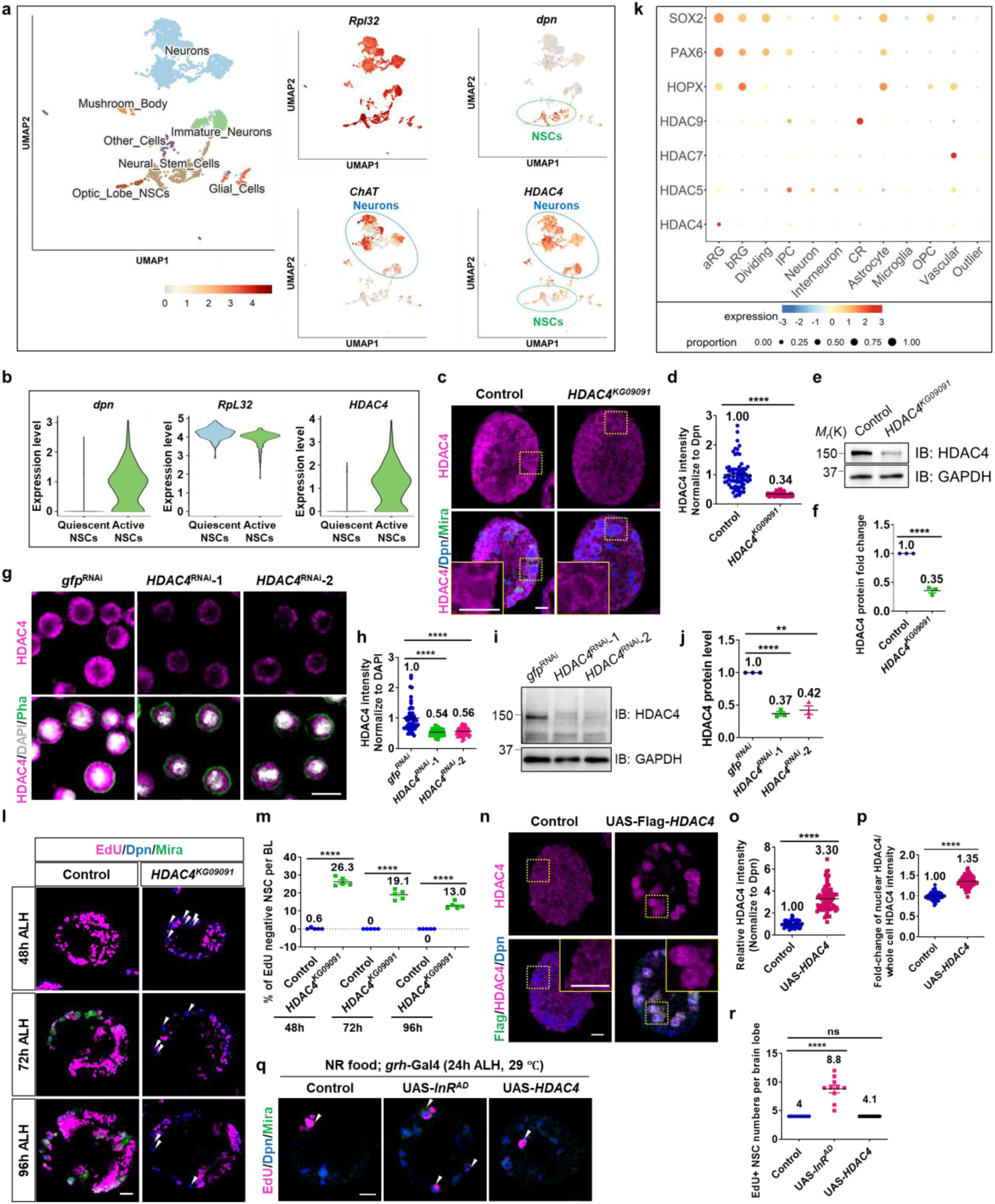
HDAC4 is expressed in the larval brain and necessary for NSC reactivation. **a, b** Analysis of the single cell RNA-sequencing dataset from late first instar larvae at 16 h ALH^67^, *RpL32* works as a house keeping gene, *dpn* is an NSC marker here works for a positive control; ChAT is a neuron marker. **c** Larval brain lobes from control (*yw*) and *HDAC4^KG09091^* at 24 h ALH. **d** Quantification graph of HDAC4 intensity (normalized to Dpn) in NSCs in (**c**). **e** Fly larval brains from yw control and *HDAC4^KG09091^* at 24 h ALH were dissected to extract proteins for western blot. **f** Quantification of relative HDAC4 protein levels in (**e**). **g, i** S2 cells were treated with double strand RNA against *gfp* or *HDAC4*. **g** Cells were labeled with anti-HDAC4 antibody, Phalloidin and DAPI. **i** Cells were harvested for Western blot. **h** Quantification graph of HDAC4 intensity (normalized to DAPI) in (**g**). **j** Quantification graph of HDAC4 protein levels (normalized to GAPDH) in (**i**). **k** Analysis of the single cell RNA-sequencing dataset from human fetal brain ^69^, human class IIa HDACs are highly expressed in human fetal brain. HDAC4 expresses in human NSCs: apical radial glial cells (aRGCs). SOX2, an NSC marker; PAX6, an apical radial glia cells marker; HOPX, a basal radial glia cells marker. **l** At 48h, 72h and 96h ALH, larval brain lobes from control (yw) and *HDAC4^KG09091^* were analyzed for EdU incorporation. NSCs were marked by Dpn and Mira. White arrows point to EdU negative quiescent NSCs. **m** Quantification graph of EdU- NSCs per BL for genotypes in (**l**). **n** Larval brain lobes in control (β-galRNAi) and UAS-*HDAC4* driven by *grh-*Gal4 at 24 h ALH. **o** Quantification graph of HDAC4 intensity (normalized to Dpn) in NSCs in (**n**). **p** At 24 h ALH, larval brain lobes from control (β-galRNAi), UAS-*InR^AD^* (positive control) and UAS-*HDAC4* driven by *grh-*Gal4 were analyzed for EdU incorporation. **q** Quantification graph of EdU+ NSCs number per BL for genotypes in (**p**). Data are presented as mean ± SD. ns for P > 0.05, **** for P ≤ 0.0001. Scale bars: 10 μm.

**Supplementary Figure 2.**
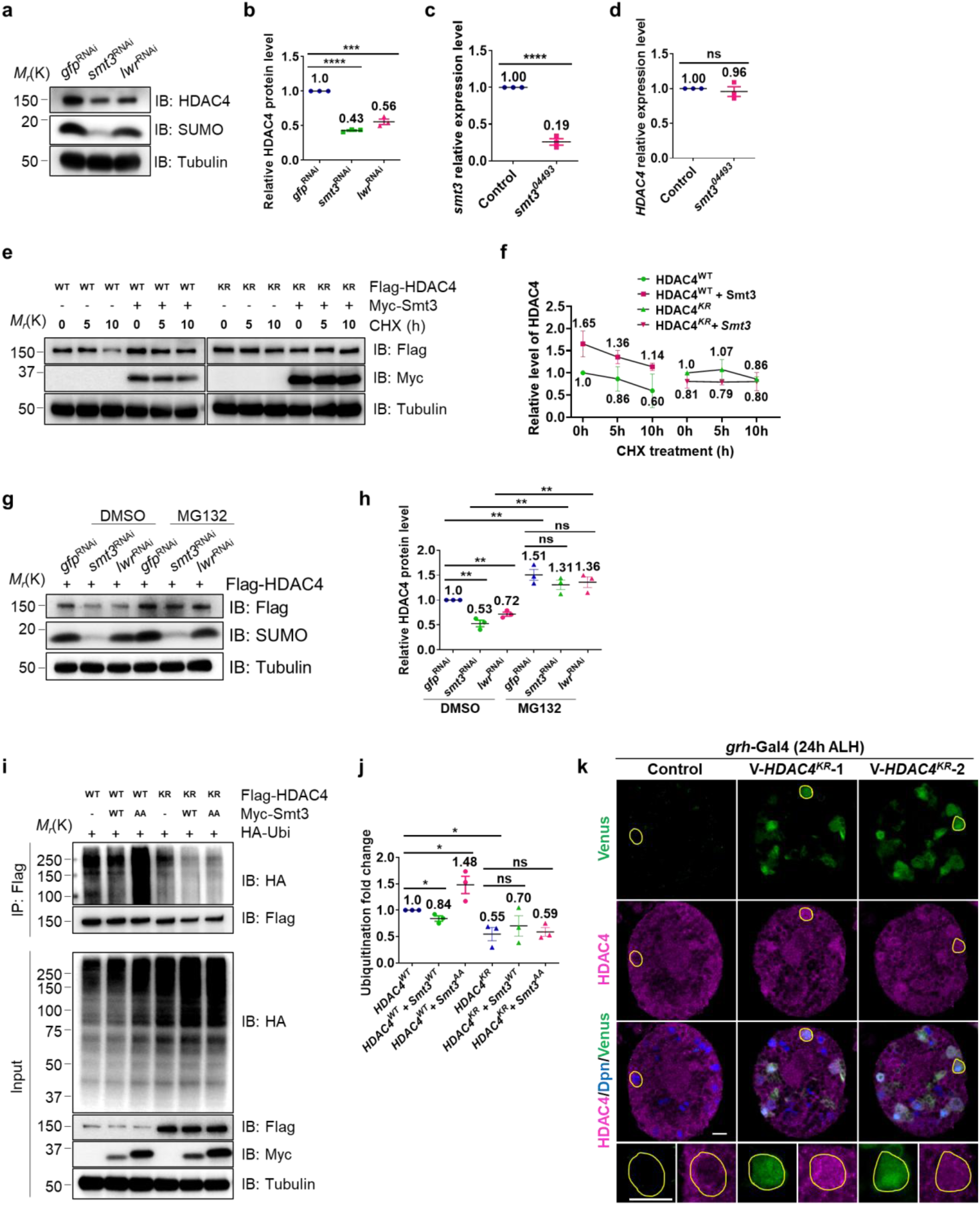
HDAC4 SUMOylation stabilizes HDAC4 protein by inhibiting its ubiquitin-proteasome degradation. **a** S2 cells were treated with double stranded RNA against *gfp*, *smt3* or *lwr* for 72 h to induce gene knockdown. Proteins were extracted for Western blot. **b** Quantification graph of HDAC4 protein levels (normalized to Tubulin) in (**a**), n=3. Control (gfpRNAi), 1; smt3RNAi, 0.43 ± 0.02, P = 8.8E-07; lwrRNAi, 0.56 ± 0.07, P = 0.0004. **c, d** Quantification of *smt3* and *HDAC4* mRNA fold enrichment in real-time qPCR assay. At 24 h ALH, larva brains from yw control and *smt3^04493^* were dissected for qPCR, n = 3. **e** S2 cells were treated with cycloheximide (CHX) for indicated intervals and collected for Western blotting with indicated antibodies. **f** Quantification graph of HDAC4 protein levels (normalized to Tubulin) in (**e**), n=3. Flag-HDAC4, 0 h, 1; 5 h, 0.86 ± 0.28; 10 h, 0.6 ± 0.38; Flag-HDAC4+Myc-Smt3, 0 h, 1.65± 0.3; 5 h, 1.36 ± 0.15; 10 h, 1.14 ± 0.08. Flag-HDAC4KR, 0 h, 1; 5 h, 1.07 ± 0.23; 10 h, 0.86 ± 0.06; Flag- HDAC4KR+Myc-Smt3, 0 h, 0.81 ± 0.15; 5 h, 0.79 ± 0.06; 10 h, 0.8 ± 0.2. **g** S2 cells were transfected with Flag-HDAC4 and treated with double stranded RNA against *gfp*, *smt3* or *lwr* for 72 h to induce gene knockdown. 4 h before harvest, S2 cells were treated with DMSO or MG132 as indicated, and then cells were collected for Western blotting with indicated antibodies. **h** Quantification graph of HDAC4 protein levels (normalized to Tubulin) in (**g**), n=3. DMSO control group, gfpRNAi, 1; smt3RNAi, 0.53 ± 0.11, P = 0.002; lwrRNAi, 0.72 ± 0.07, P = 0.002; MG132 group, gfpRNAi, 1.51± 0.19; smt3RNAi, 1.31 ± 0.16, P = 0.24; lwrRNAi, 1.36 ± 0.19, P = 0.39; **i** Ubiquitination assay in S2 cells, cells were transfected with indicated plasmids. S2 cells were treated with 10 µM MG132 for 4 h before harvest. IP was conducted with anti-Flag antibody. Western blots were performed by using anti-HA and anti-Flag to detect ubiquitinated HDAC4 and overall levels of Flag-HDAC4, respectively. **j** Quantification of HDAC4 ubiquitination for (**i**). HDAC4WT, 1; HDAC4WT + Smt3WT, 0.84 ± 0.09, P = 0.03; HDAC4WT + Smt3AA, 1.48 ± 0.28, P = 0.04; HDAC4KR, 0.55 ± 0.22, P = 0.02; HDAC4 KR + Smt3WT, 0.7 ± 0.33, P = 0.53; HDAC4 KR + Smt3AA, 0.59 ± 0.14, P = 0.8. **k** Re-analysis of a published dataset of single cell RNA-sequencing obtained from late first instar larvae at 16 h ALH^67^ for the expression of SUMO and Ubiquitin in quiescent NSCs, pre-active NSCs and reactivating NSCs. **l** At 24 h ALH, overexpression of Venus-*HDAC4^K902R^* in NSCs, Venus labels the overexpressed HDAC4, HDAC4 labels the endogenous HDAC4. Data are presented as mean ± SD. ns for P > 0.05, * for 0.05≤ P ≤ 0.01, ** for P ≤ 0.01, *** for P ≤ 0.001, **** for P ≤ 0.0001. Scale bars: 10 μm.

**Supplementary Figure 3.**
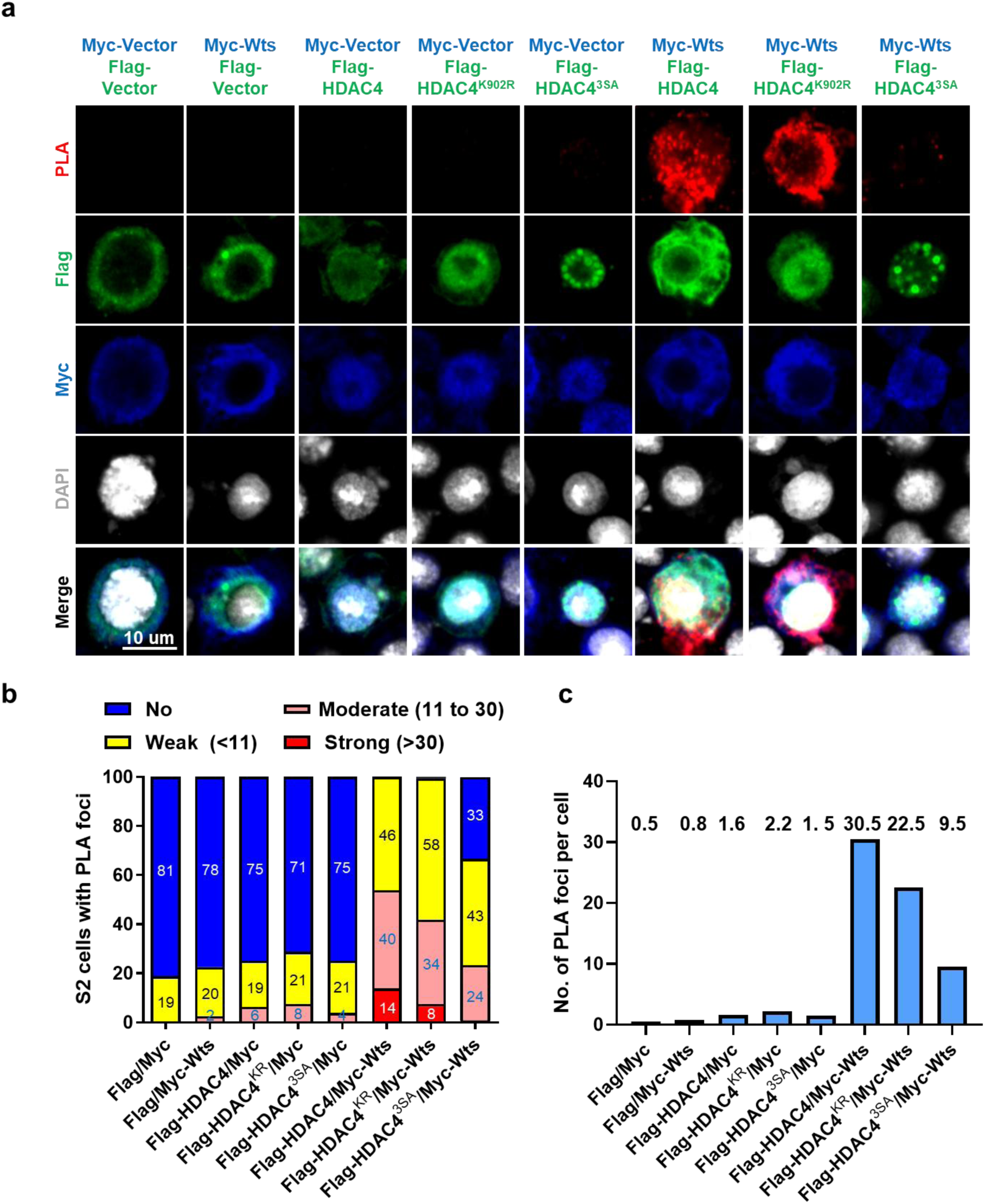
Elevated nuclear HDAC4 localization disrupts its association with Wts. **a** *In situ* PLA assay between Myc-Wts and Flag-HDAC4 (WT/KR/3SA) in S2 cells. S2 cells transfected with two of the indicated plasmids were stained for Flag, Myc, and DNA and detected for PLA signal. **b** Quantification graphs showing the percentage of S2 cells with no PLA signal, weak (<11 foci), moderate (11–30 foci), and strong (>30 foci) PLA signals for (**a**). (**c**) Quantification graph of the average number of PLA foci per cell in (**a**). Scale bars: 10 μm.

**Supplementary Figure 4.**
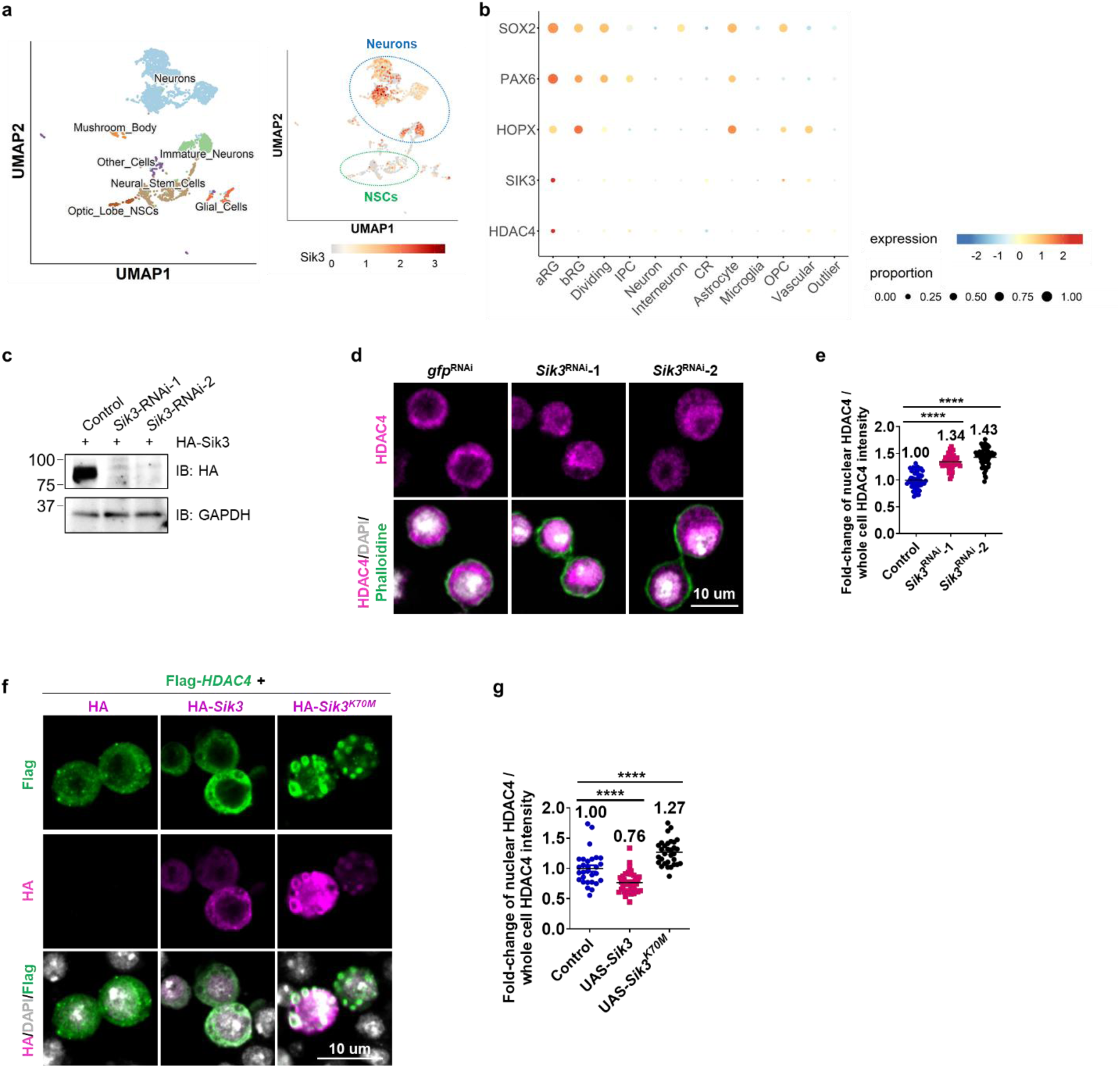
SIK3 is highly expressed in brains and results in HDAC4 cytoplasmic localization. **a** Re-analysis of a published dataset of single cell RNA-sequencing obtained from late first instar larvae at 16 h ALH^67^ for the expression of *Sik3* in brains. **b** Re-analysis of the single cell RNA-sequencing dataset from human fetal brain ^69^, *Sik3* expressed in human NSCs: apical radial glial cells (aRGCs). SOX2 is an NSC marker; PAX6 is an apical radial glia cell marker; HOPX is a basal radial glia cell marker. **c** S2 cells were transfected with HA-Sik3 and treated with dsRNA against *gfp* or *Sik3* for 72 h. Cells were harvested for Western blot to test the knockdown efficiency of *Sik3*. **d** S2 cells were treated with dsRNA against *gfp* or *Sik3* for 72 h. Cells were labeled with anti-HDAC4 antibody, Phalloidin and DAPI. **e** Quantification graph of the fold-change of nuclear HDAC4 intensity/whole cell HDAC4 intensity in (**d**). **f** S2 cells were transfected with the indicated constructs. Cells were labeled with anti-Flag and anti-HA antibodies and DAPI. **g** Quantification graph of the fold-change of nuclear HDAC4 intensity/whole cell HDAC4 intensity in (**f**). Data are presented as mean ± SD. **** for P ≤ 0.0001. Scale bars: 10 μm.

**Supplementary Figure 5.**
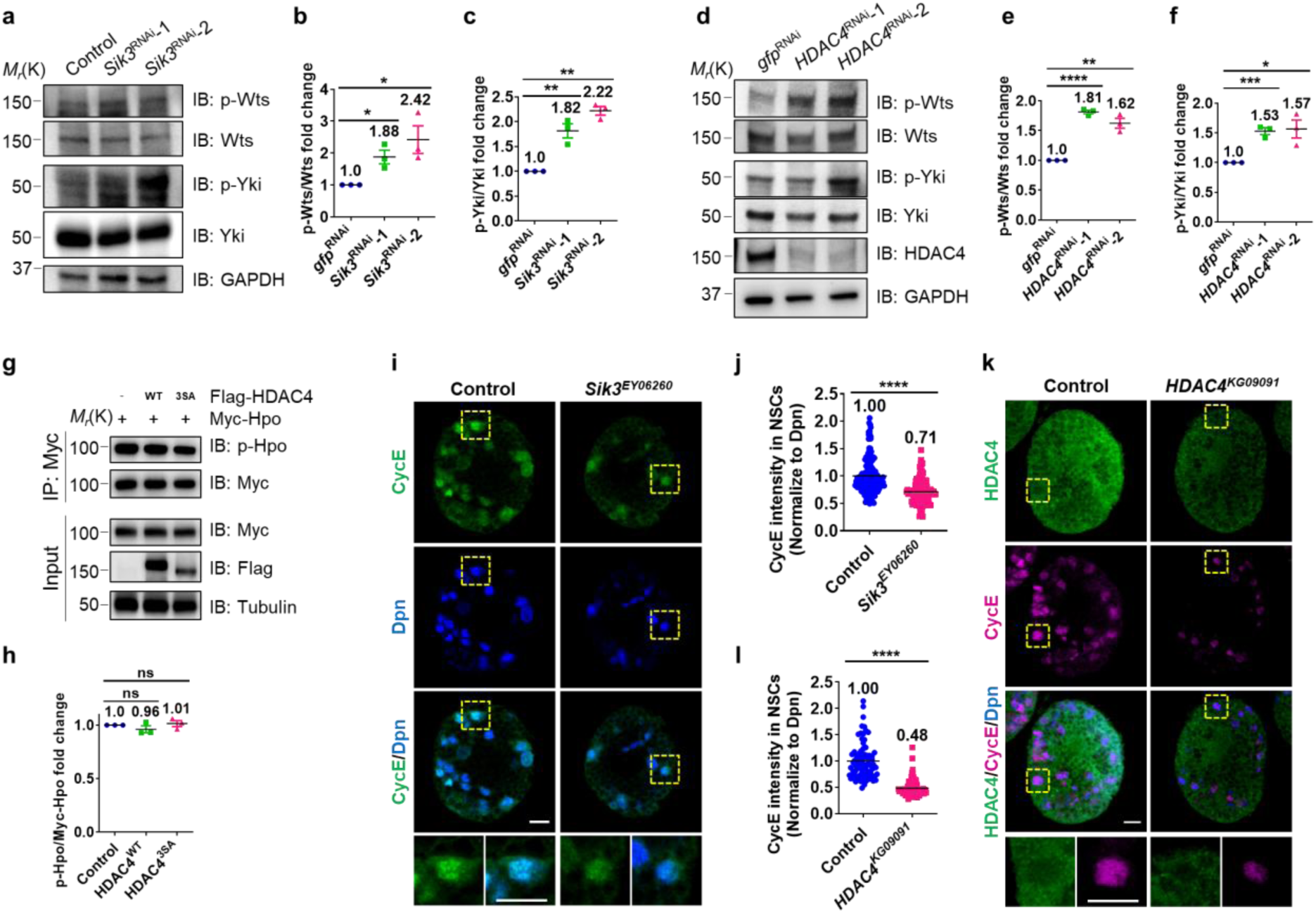
SIK3-HDAC4 promotes the expression of Hippo pathway target genes in NSCs. **a, d** S2 cells were treated with dsRNA against *gfp* or *Sik3* or *HDAC4* for 72 h. Cells were then harvested for Western blot. **b, e** Quantifications for (**a, d**), respectively. **b** *gfp*RNAi, 1; *Sik3*RNAi-1, 1.88 ± 0.37, P = 0,015; *Sik3*RNAi-2, 2.42 ± 0.75, P = 0.03. **e** *gfp*RNAi, 1; *HDAC4*RNAi-1, 1.81 ± 0.05, P = 1.4E-05; *HDAC4*RNAi-2, 1.62 ± 0.14, P = 0.0017. **c, f** Quantifications for (**a, d**), respectively. **c** *gfp*RNAi, 1; *Sik3*RNAi-1, 1.82 ± 0.25, P = 0.0046; *Sik3*RNAi-2, 2.22 ± 0.15, P = 0.00015. **f** *gfp*RNAi, 1; *HDAC4*RNAi-1, 1.53 ± 0.1, P = 0.0008; *HDAC4*RNAi-2, 1.57 ± 0.26, P = 0.02. **g** S2 cells were transfected with the indicated constructs, then harvested for IP and Western blot for p-Hpo T195. **h** Quantifications for (**g**). Control, 1; *HDAC4*, 0.96 ± 0.06, P = 0.3; *HDAC43SA*, 1.01 ± 0.05, P = 0.6. **i, k** Larval brain lobes from control (*yw*), *Sik3^EY06260^*or *HDAC4^KG09091^* at 24 h ALH been labeled with CycE and Dpn. **j, l** Quantification graph of CycE intensity (normalized to Dpn) in NSCs in (**I, k**). **j** Control (yw), 1 ± 0.2, n = 10; *Sik3^EY06260^*, 0.61 ± 0.2, P = 3.6E-07, n = 10. **l** Control (yw), 1 ± 0.34, n = 11; *HDAC4^KG09091^*, 0.48 ± 0.16, P = 3.1E-27, n = 11. Data are presented as mean ± SD. ns for P > 0.05, * for 0.05≤ P ≤ 0.01, ** for P ≤ 0.01, *** for P ≤ 0.001, **** for P ≤ 0.0001. Scale bars: 10 μm.

**Supplementary Figure 6.**
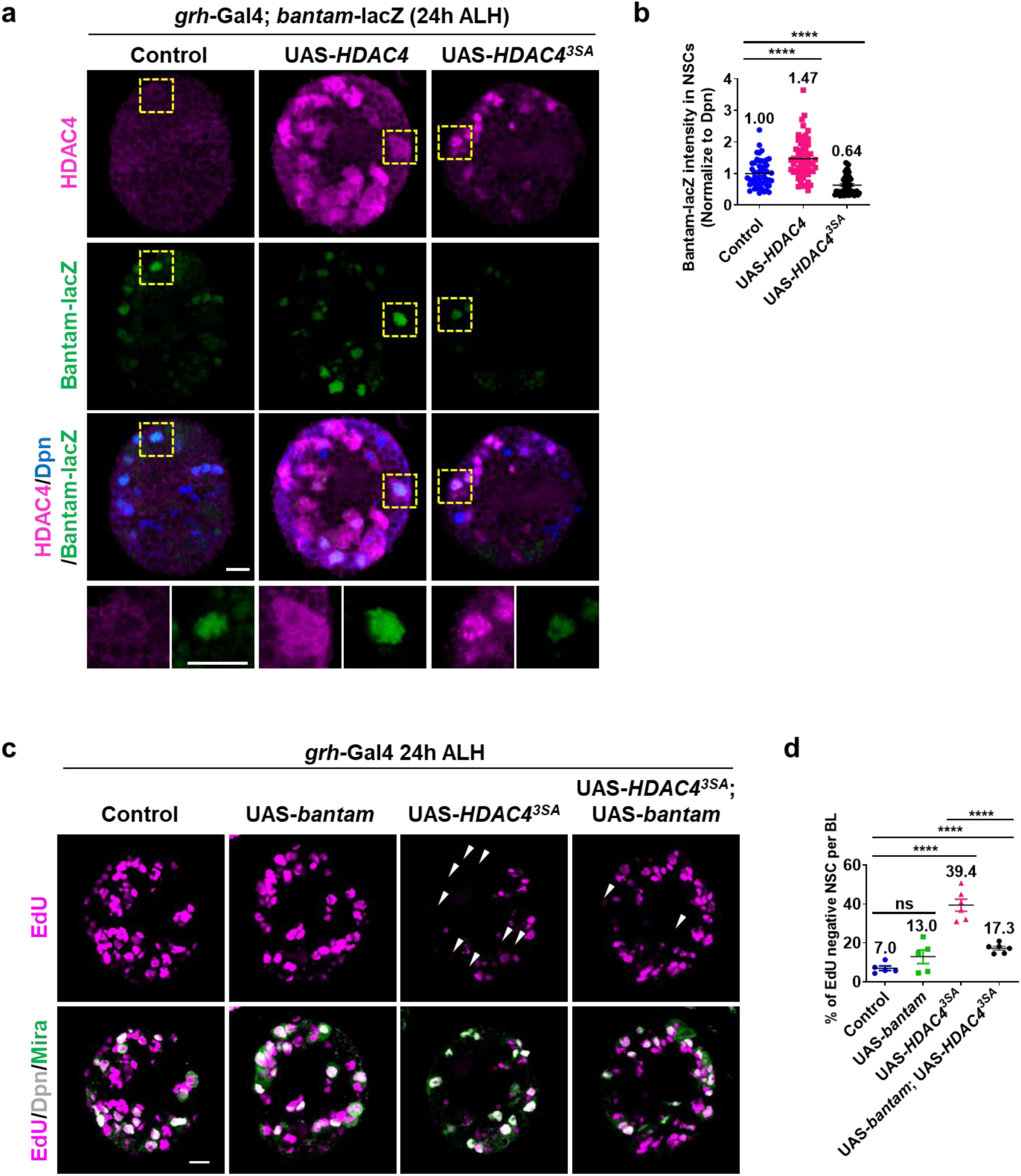
HDAC4 promotes the expression of *bantam* in NSCs. **a** Larval brain lobes with the indicated genotypes at 24 h ALH been labeled with HDAC4, Bantam-lacZ and Dpn. **b** Quantification graph of Bantam-lacZ intensity (normalized to Dpn) in NSCs in (**a**). Control (β-galRNAi), 1 ± 0.44, n = 10; UAS-*HDAC4*, 1.47 ± 0.61, P = 1.8E-05, n = 10; UAS-*HDAC43SA*, 0.64 ± 0.31, P = 8.2E-06, n = 10. **c** At 24 h ALH, larval brain lobes from various genotypes were analyzed for EdU incorporation. NSCs were marked by Dpn and Mira. White arrows point to EdU negative qNSCs. **d** Quantification of EdU- NSCs per BL for (**c**). Control (β-galRNAi), 7 ± 2.7, n = 5; *UAS-bantam*, 13 ± 7.9, P = 0.15, n = 5; UAS-*HDAC43SA*, 39.4 ± 7.2, P = 8.8E-06, n = 6. *UAS-bantam*; UAS-*HDAC43SA*, 17.3 ± 2.7, P = 9.4E-05, P = 4.9E-05, n = 6. Data are presented as mean ± SD. ns for P > 0.05, **** for P ≤ 0.0001. Scale bars: 10 μm.

**Supplementary Table 1.** Classification of Histone Deacetylases in human and *Drosophila*.

| Classification |  | Vertebrates | *Drosophila* |
| --- | --- | --- | --- |
| Class I |  | HDAC1 | HDAC1 |
|  |  | HDAC2 |  |
|  |  | HDAC3 | HDAC3 |
|  |  | HDAC8 |  |
| Class II | IIa | HDAC4 | HDAC4 |
|  |  | HDAC5 |  |
|  |  | HDAC7 |  |
|  |  | HDAC9 |  |
|  | IIb | HDAC6 | HDAC6 |
|  |  | HDAC10 |  |
| Class IV |  | HDAC11 | HDAC11 |
| Class III |  | SIRT1 | Sirt1 |
|  |  | SIRT2 | Sirt2 |
|  |  | SIRT3 |  |
|  |  | SIRT4 | Sirt4 |
|  |  | SIRT5 |  |
|  |  | SIRT6 | Sirt6 |
|  |  | SIRT7 | Sirt7 |

**Supplementary Table 2.**
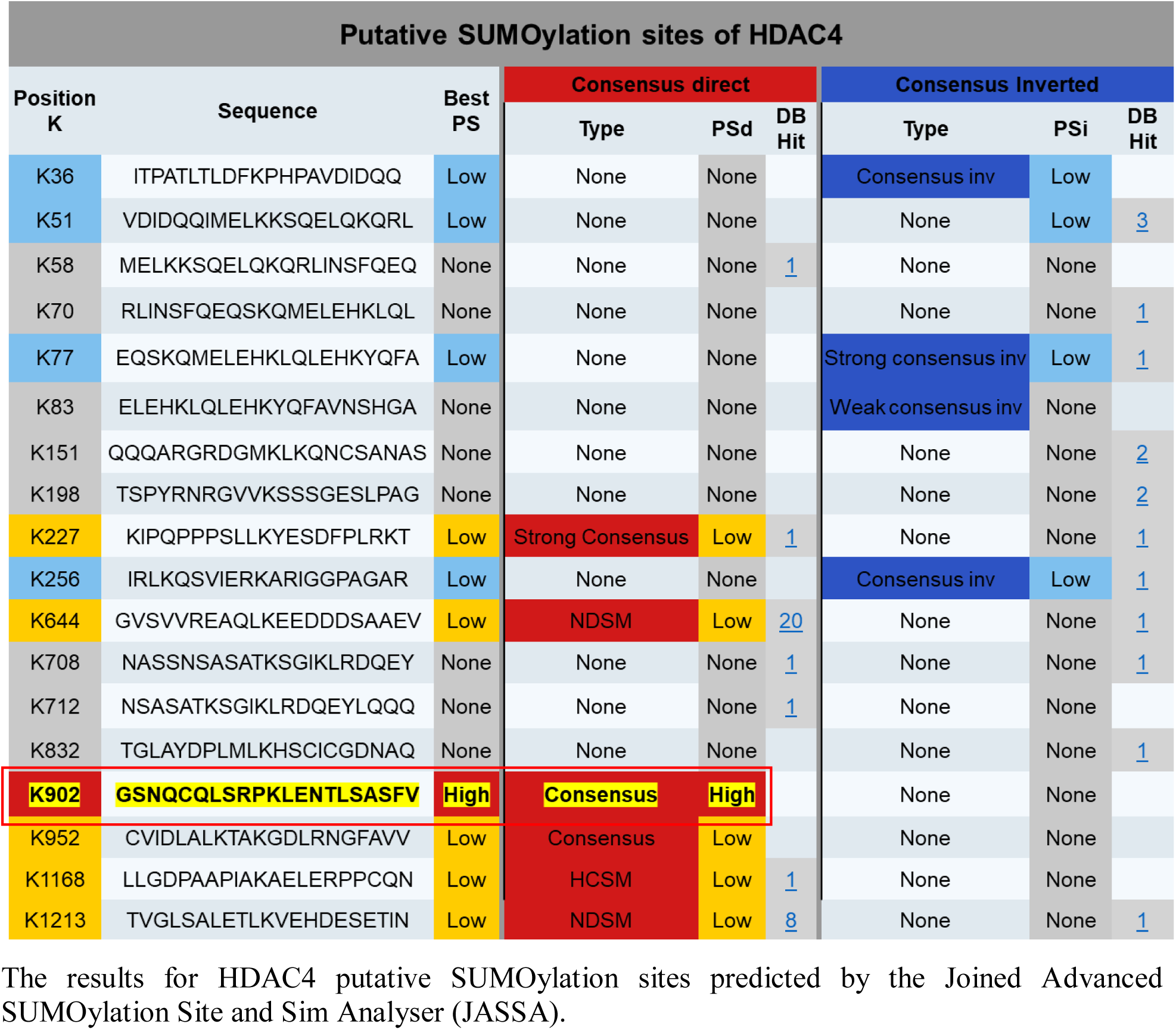
HDAC4 K902 is predicted to be SUMOylated.

**Supplementary Table 3.**
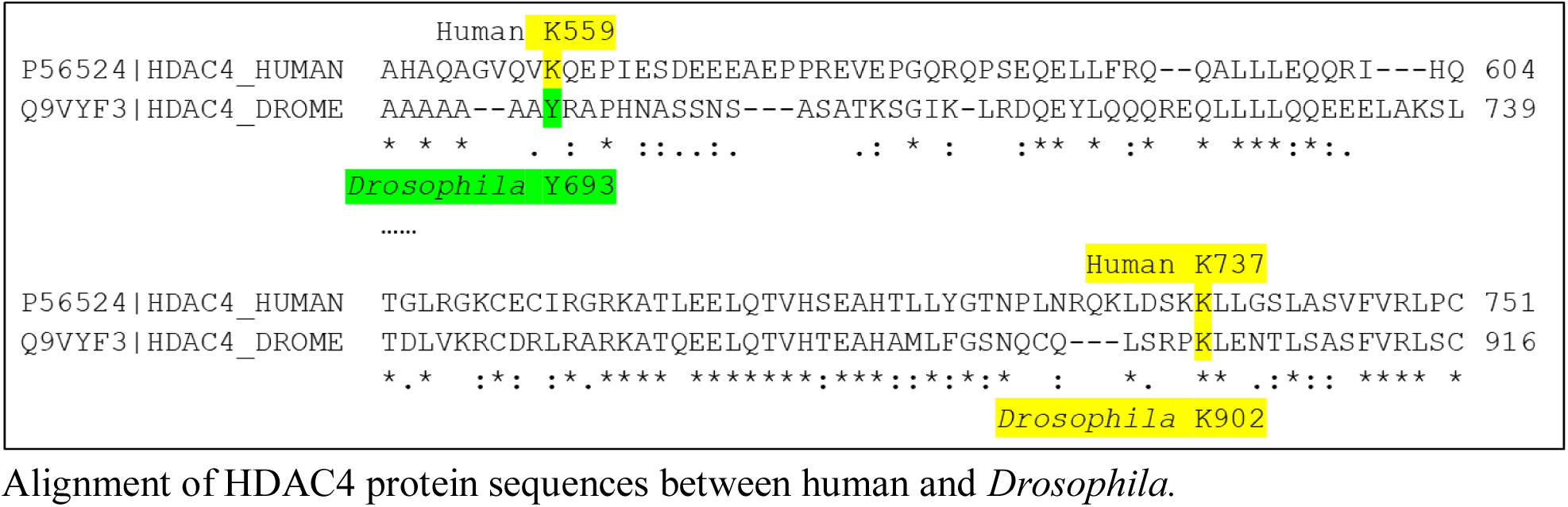
HDAC4 K902 is not equivalent to human K559.

**Supplementary Table 4.**
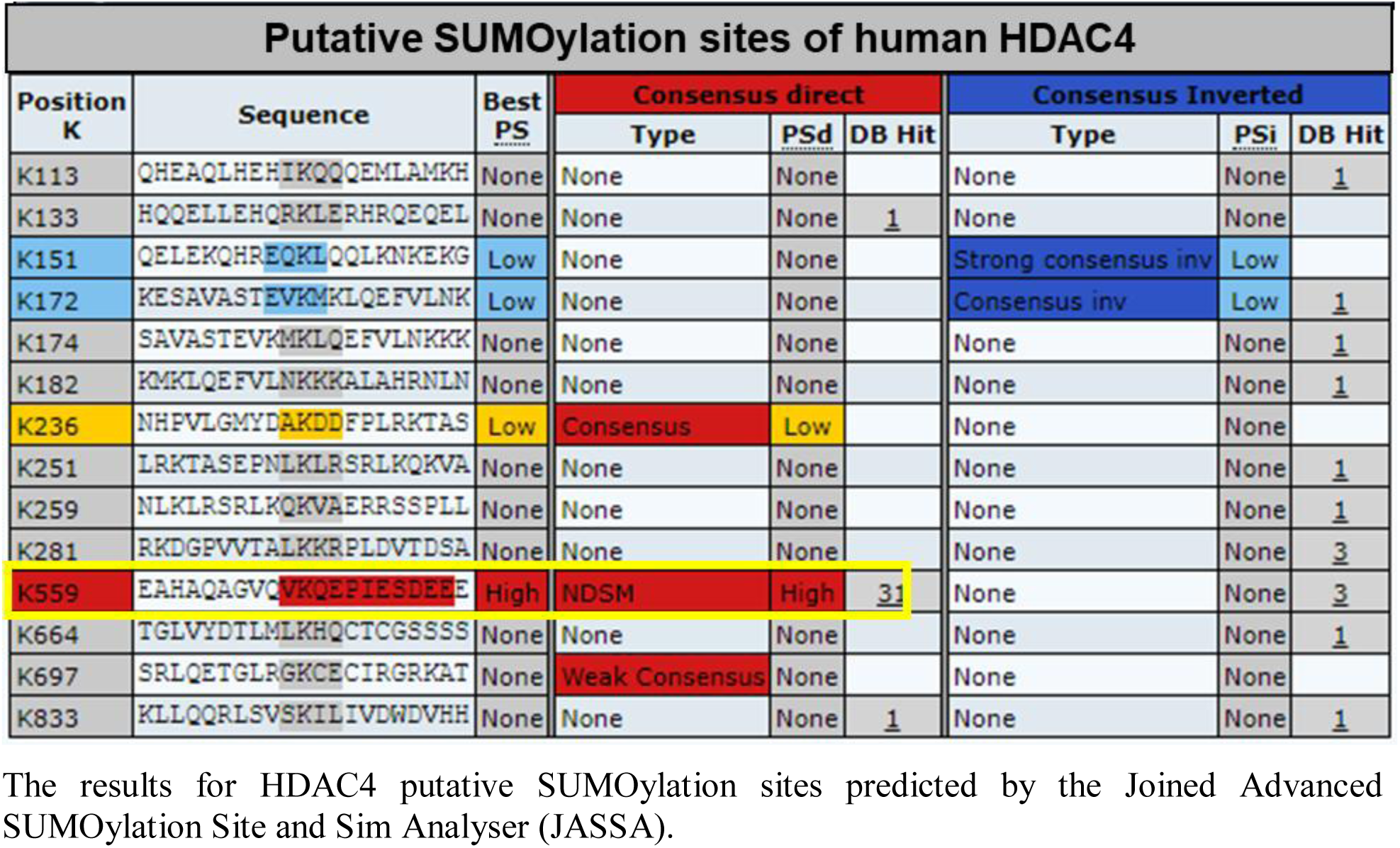
Human HDAC4 K559 is predicted to be SUMOylated but not K737.

**Supplementary Table 5.**
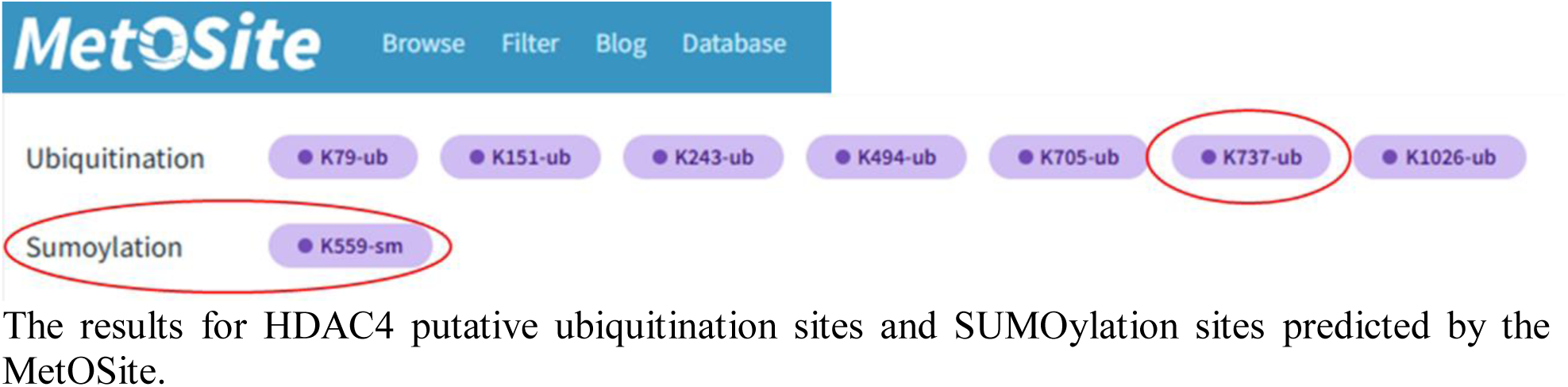
Human HDAC4 K737 is predicted to be ubiquitinated but not SUMOylated.

